# *In Ovo* Sexing and Genotyping using PCR techniques: A Contribution to the 3R Principles in Chicken Breeding

**DOI:** 10.1101/2025.11.11.687802

**Authors:** C. Dierks, A. Förster, D. Meunier, R. Preisinger, C. Klein, S. Weigend, S. Altgilbers

**Affiliations:** Institute for Laboratory Animal Science and Central Animal Facility, Hannover Medical School, Hannover, Germany; EW GROUP GmbH, 49429, Visbek, Germany; National Avian Research Facility, The Roslin Institute and Royal (Dick) School of Veterinary Studies, University of Edinburgh, United Kingdom; Friedrich-Loeffler-Institut – Federal Research Institute for Animal Health (FLI), Institute of Farm Animal Genetics, Germany

**Keywords:** *in ovo* sexing, *in ovo* genotyping, KASP, 3R principles

## Abstract

Early sex determination and genotyping of chicken embryos is crucial for ethical and resource-efficient animal research, addressing concerns about surplus animals. We developed a reproducible workflow using whole genome amplification combined with Kompetitive Allele Specific PCR (KASP) and standard endpoint PCR to perform *in ovo* sexing and genotyping from embryonic day four (ED4/96h) onwards. The overall efficiency improved with embryonic age. Both standard PCR and KASP provided high success for sexing and genotyping (70–100% of samples yielding a result) and accuracy (92–100%) across multiple chicken lines, including a genetically modified line. Optimal reliability and hatchability were achieved when sampling at ED7. These PCR-based techniques enable precise early identification of sex and genotype, allowing selective removal of unwanted embryonated eggs before the assumed onset of nociception. This approach supports more humane and efficient practices in chicken breeding and research, contributing to the replacement, reduction and refinement (3R) principles in animal experimentation.

## Introduction

The chicken has emerged as a popular and valuable animal model, contributing significantly to numerous discoveries in biomedical research^1^. It is particularly well-suited for studying early embryo development, immunology, toxicology, oncology, conservation biology, virology disease mechanisms, and epigenetics^2–6^. The fact that it can be accessed without harming the hen is a major advantage of this model^7^. Although the chicken embryo has often been considered a partial replacement model to mice^8,9^, it is crucial to recognize that chicken embryos may also experience pain^10–12^. This has sparked ethical and political debates, particularly concerning the layer industry’s practice of culling male chicks shortly after hatching, for economic reasons. In response, there has been growing interest in finding alternatives to this practice^13^. Research on nociception and pain perception has contributed to the development of early *in ovo* sexing techniques and the identification of optimal developmental stages for their application^14,15^.

Considering the capacity for pain perception of chicken embryos prior to hatching, several countries have implemented regulations on the destruction of embryonated eggs and the culling of chicks^16^.

In France, the culling of hatched chicks from *Gallus gallus* lines intended for the production of eggs for human consumption is prohibited (R214-17, R214-78 Code rural et de la pêche maritime)^17,18^, but exemptions apply: chicks intended for animal food production and incorrectly sexed chicks may be culled^16–18^.

In the United Kingdom, chicken embryos are not classified as mature vertebrates under the Animals (Scientific Procedures) Act 1986 (ASPA) until day 15 of incubation (embryonic day (ED) 15), which constitutes two-thirds of the incubation period^9,19^. This means that manipulating or culling chick embryos can take place outwith ASPA licensing up to ED14. Germany has adopted a distinct approach, driven by the latest findings on pain perception in chicken embryos^10–12^. It bans any procedure, including terminating incubation, that could result in the death of a chicken embryo after ED12. However, this restriction on disposing of embryonated eggs only applies if *in ovo* sexing was performed (section 4c German Animal Welfare Act (Tierschutzgesetz))^20^. If *in ovo* sexing was not conducted, the general ban on culling chicks still applies post-hatch.

Crucially, chicken embryos used for research purposes are not yet protected, unless *in ovo* sexing is performed. The German Animal Welfare Regulation Governing Experimental Animals (Tierschutz-Versuchstierverordnung) only mandates protection for mammalian fetuses from the last third of their normal development before birth^21^. Consequently, procedures involving chicken embryos, such as microinjections, only require approval and notification if the chicks are intended to hatch.

The development of specialized laboratory strains and specific pathogen-free lines has further enhanced the chicken’s utility as a research model^22–25^. Genetically altered lines, such as the iCaspase9 surrogate host chicken^23^, offer promising avenues for reducing the number of animals to address research questions^26^. Careful planning and breeding strategies are crucial to minimize the number of animals required for experiments^27^. Some hatched chicken may not possess the desired traits or sex and can only be used partially in experiments, for example as control animals or for further breeding^28^. Culling an animal without a valid scientific or ethical justification is prohibited by section 17 (1) of the German Animal Welfare Act. The UK Guidance on the operation of ASPA requires researchers to strictly adhere to the replacement, reduction and refinement (3R) principles and use breeding management to minimize surplus animals. In contrast, German and Austrian Animal Welfare Law contain the unique term ‘reasonable cause’ for culling, demanding legal justification for the killing of any animal. While scientific research is generally a ‘reasonable cause’, this justification is not automatically granted for surplus animals that were never used in an experiment. This distinction fuels the ongoing ethical debate and calls into question reduction strategies such as cascade utilization^29^. *In ovo* sex determination techniques, particularly molecular methods such as Polymerase Chain Reaction (PCR) or hormone analysis, enable early identification of an embryo’s sex or other genetical characteristics from embryonic tissue collected during incubation, including pieces from the chorioallantoic membrane (CAM) or amniotic fluid^30–40^. Recent advances have enabled minimally invasive puncture of the eggshell to collect allantoic fluid, followed by PCR analysis^41–43^.

The objective of this study was to develop a reliable and user-friendly method for *in ovo* sexing and genotyping of chicken lines used for research purposes. Early *in ovo* sex determination, which does not require complex equipment, would benefit breeding management, particularly for genetically engineered (GE) chicken lines. This capability is crucial for all laboratories committed to the 3Rs principles as it directly enhances animal welfare by allowing the removal of surplus embryos before the assumed onset of nociception. By providing a robust screening tool, the method mitigates significant issues concerning biosecurity, compliant waste disposal, and excessive animal resource demands. While beneficial globally, this solution is particularly vital for new transgenesis laboratories, especially those in Low- and Middle-Income Countries (LMICs), which often rely on less-efficient initial breeding methods due to a lack of surrogate host systems^23,44,45^.

## Materials and Methods

### Animal Experiments and Animal Care/Ethical statements

The study is reported in accordance with the ARRIVE guidelines^46,47^. Sexing chickens before hatch is a common practice in hatcheries in Germany. In the present study, samples from chicken embryos were obtained prior to the assumed onset of nociception (ED13)^10–12^; this did not require an animal experiment application, which is in accordance with the legal framework for animal experiments in Germany (German Animal Welfare Act (TierSchG)^20^ and Animal Welfare Regulation Governing Experimental Animals (TierSchVerV)^21^). The German legislation is based on the overarching EU directive 2010/63/EU. Allantoic puncture and blood sample collection from hatched chicks during the Phase I study were approved by the Animal Care and Use Committee of Lower Saxony (Niedersächsisches Landesamt für Verbraucherschutz und Lebensmittelsicherheit), Oldenburg, Germany (approval number: AZ 33.9-42502-05-10A064)).

### Experimental design

The study consisted of four parts to evaluate the feasibility and impact of the *in ovo* sexing and genotyping techniques.

The Pilot study aimed to evaluate the feasibility of the techniques at various stages of embryonic development.

The Phase I study was conducted on a larger number of eggs and focused on the impact of *in ovo* sexing and genotyping on the hatchability of the sampled eggs. Both Pilot and Phase I studies were part of the EU Horizon 2020 project Innovative Management of Animal Genetic Resources (IMAGE; https://www.imageh2020.eu/), utilizing marker-assisted introgression to introduce a single monogenic dominant trait (blue egg shell color) from a gene bank population into a high performing white-egg layer chicken line^48,49^.

The Phase II study focused on *in ovo* sexing at early embryonic stages and evaluation of subsequent hatch rates utilizing commercial white-egg and brown-egg layer chicken lines.

The Phase III study was a proof-of-concept study to determine the feasibility of applying these techniques (sexing and genotyping) to a genetically modified chicken line (iCaspase9 surrogate host)^23^ for maintenance breeding.

Experimental time points for fluid collection are detailed in Fig. 1. To guide the selection of appropriate time points for allantoic puncture, an *ex ovo* culture was first set up to assess the spatial extent of the CAM and allantoic sac across different incubation days. Early *in ovo* sampling confirmed that various internal egg fluids contained DNA. To reflect this, the term “*in ovo* fluid” is used for all samples collected before ED7. From ED7 onward, we are confident that the samples represent allantoic fluid.

**Fig. 1.**
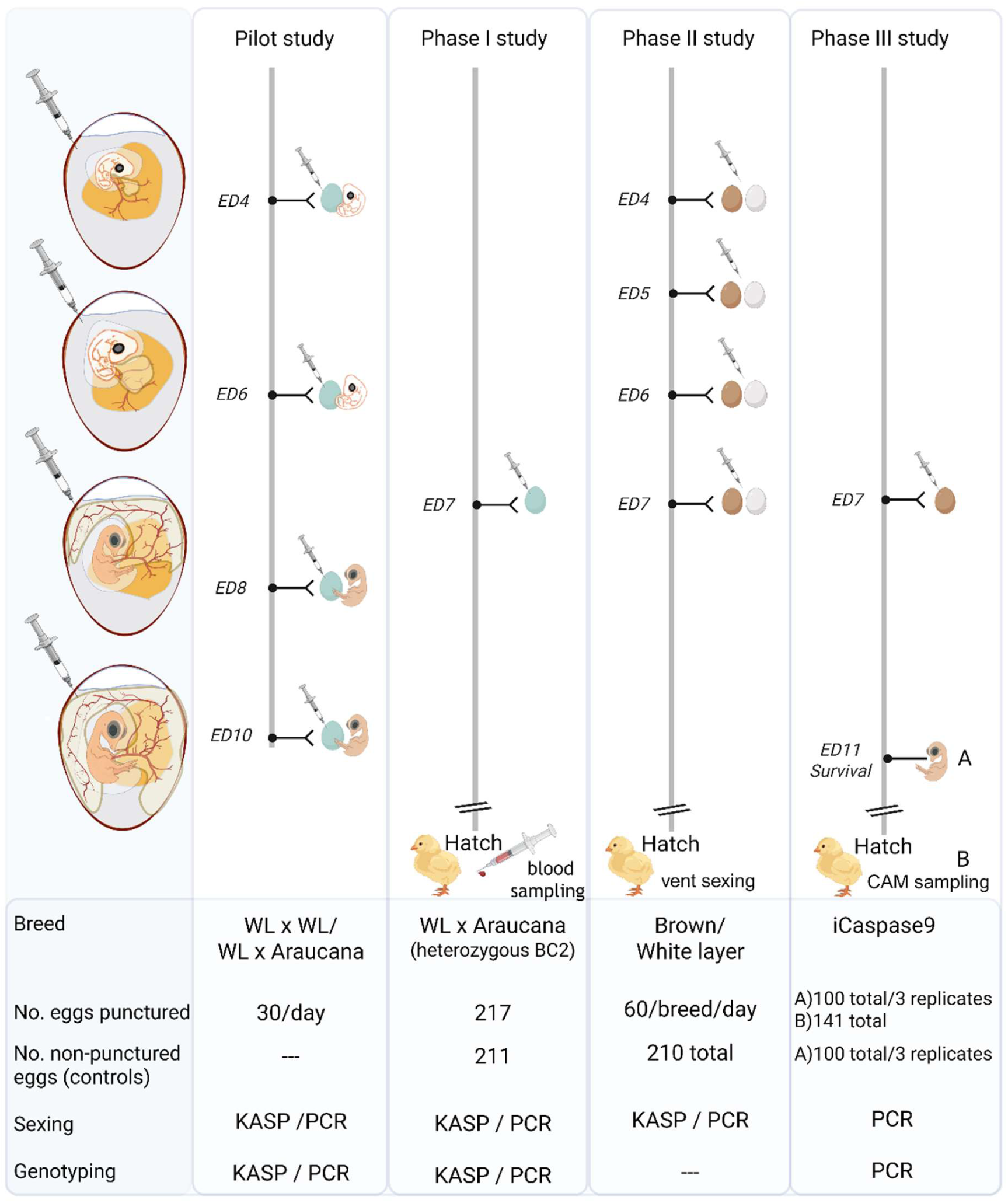
Experimental Workflow for *in ovo* or Allantoic Fluid Collection from Embryonated Eggs. Schematic representation of chicken embryo development from day five (ED4) to eleven (ED10) of incubation, highlighting rapid allantoic vesicle formation and chorioallantoic membrane (CAM) development. The study comprised a Pilot study and three subsequent studies (Phase I, II, III). In each study, eggs were punctured at various time points during incubation to obtain fluid samples for DNA extraction and PCR analysis. In the Pilot study, embryos were dissected after puncture to obtain tissue samples for sex verification. In Phase I and Phase II studies, embryos were allowed to hatch after sampling, and blood samples were collected or vent sexing was performed to verify *in ovo* sexing accuracy. In the Phase III study, eggs from genetically modified embryos were punctured at ED7 and then returned to the incubator until ED11. A) Survival rates were assessed at ED11 post-puncture, and a tissue sample was collected from each embryo. B) From 141 fertilized eggs punctured on ED7, a defined subset was allowed to hatch, and CAM samples were collected for DNA extraction. (Created in BioRender. Altgilbers, S. (2025) https://BioRender.com/gy26oy3)

### *Ex ovo* culture

A shell-less culture protocol was used from ED2 to ED11 as previously described^50,51^. *Ex ovo* culture was initiated with embryos at Hamburger and Hamilton^52^ (HH) stage 15-17, corresponding to approximately 65 hours of incubation. Embryos in weigh boats were covered with a petri dish lid and incubated without rotation in a fully automated digital incubator (Ova-Easy 100 Advance Series II Cabinet Incubator, BRINSEA PRODUCTS LTD, North Somerset, UK) at 65 % relative humidity using a humidity pump (Ova-Easy/TLC Advance Humidity Pump, BRINSEA PRODUCTS LTD, North Somerset, UK). Photographs of embryos and gonads, and allantoic cavity measurements (96-144h; n=10-15 embryos/incubation day) were acquired by light microscopy using an Olympus SZ61 stereomicroscope equipped with an Olympus EP50 camera and the associated EPView Windows software or iOS App (EVIDENT Europe GmbH, Hamburg, Germany). Images shown in Fig. S1 were captured using an iPhone 13 Pro (12 MP camera) and allantoic cavity measurements (168-192h) were taken manually with a ruler, and scale bars were subsequently set in ImageJ 1.48v/Java 1.6.0_20 (64-bit). Using an insulin syringe (Omnican 40; B. Braun Melsungen AG, Melsungen, Germany), food coloring was precisely injected into the allantoic or amniotic cavity under microscopic control (Olympus SZ61 stereomicroscope).

### Pilot study

White Leghorn (WL) hens were inseminated with sperm from WL or homozygous blue egg layer Araucana roosters. Eggs were stored for up to six days at 15°C. For incubation, eggs were placed in an automatic turning incubator (BRUJA, Brutmaschinen-Janeschitz GmbH, Hammelburg, Germany). The incubator was set to maintain a temperature of 37.8°C and a humidity level of 50-55 %, with eggs being turned 45° at regular intervals. Thirty fertilized eggs were incubated for a full 4 (96 hours), 6 (144 hours), 8 (192 hours), or 10 (240 hours) days. The first day of incubation was assigned as ED0. The air chamber and embryo location were identified using a LED light egg candler (Fig. 2A; Fig. S1A). The egg was positioned with the air chamber upwards (blunt end) and the puncture site was marked with a pen approximately 2-5 mm beneath the air chamber, as previously described^37^ (Fig. 2A). A small hole was drilled in the eggshell near the air chamber using an awl (Fig. 2B). Approximately 50 microliters of *in ovo* or allantoic fluids were extracted from each egg using an insulin syringe (Omnican 40; B. Braun Melsungen AG, Melsungen, Germany; Fig. 2D) and transferred to a 1.5 ml Eppendorf tube (Eppendorf SE, Germany, Hamburg; Fig. 2E). If sampling was unsuccessful after two attempts, the egg was discarded from analysis. Unfertilized eggs or early death embryos, as well as eggs with displaced air chamber were discarded (Fig. S1A-B, D). Collected samples were stored on ice and then frozen at -20°C. Following collection of *in ovo* fluid, embryos were dissected and tissue samples from all embryos were collected and frozen at -20°C for subsequent DNA extraction to confirm the results obtained from the analysis of the *in ovo* fluid samples. In addition, embryonic gonads from ED10 embryos were visually examined for sex identification (Fig. S2A-D).

**Fig. 2.**
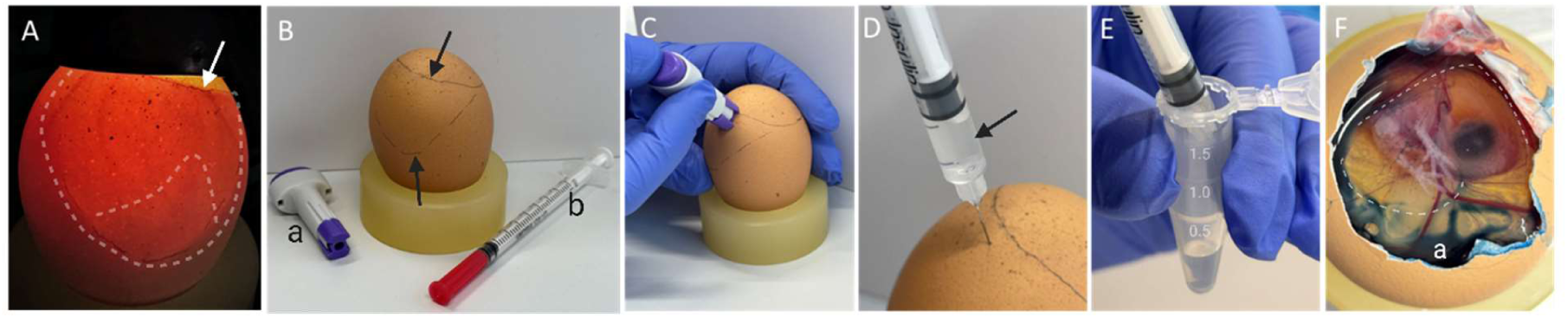
Step-by-Step Demonstration of Allantoic Fluid Collection. **(A)** Fertile brown egg (ED7) candled with an LED light. The white arrow indicates the border between egg contents and air cell at the blunt end of the egg. The white dashed lines delineate a yolk-free area of the egg, with the embryo visible at the bottom of this area, shining through the fluids within the amnion and chorioallantoic sac. **(B)** The air sac and the yolk free area of the egg, as described in **(A)**, were marked with a pencil (black arrows); **(a)** single-use lancing devices (ACCU-CHEK, Safe-T-Pro UNO, Roche AG, Basel, Switzerland), **(b)** insulin syringe (Omnican 40; B. Braun Melsungen AG, Melsungen, Germany). **(C)** A small hole was created in the eggshell using a lancing device, positioned close to the border of the air chamber. **(D)** The insulin syringe was used to aspirate the clear allantoic fluid (black arrow) through the previously created hole. For aspiration, the syringe needle was inserted to a depth of about 3-4 mm into the egg. **(E)** The aspirated allantoic fluid was transferred into a tube. **(F)** To demonstrate that the amnion was not punctured during the procedure, an egg was opened at the blunt end after the procedure. Blue food coloring **(a)** was injected through the hole in the eggshell, and no color was observed in the amnion cavity (marked with white dashed lines), confirming that only the allantoic cavity had been punctured (ED11).

### Phase I study

The hens and roosters used for egg production were obtained from the second backcross of Araucana with WL. These birds were also part of the IMAGE project. Over two generations, heterozygous roosters for the blue eggshell allele were backcrossed to WL hens. For the present study, hens that were heterozygous for the dominant blue egg-laying trait were inseminated with sperm from heterozygous roosters of the backcross generation. 480 fertile eggs were incubated in an auto-rotating incubator (PETERSIME NV, Zulte, Belgium) under standard conditions (see Pilot study). At ED7 (184h), 50 µl of fluid were extracted from half of the incubated eggs (217 total), as described in the Pilot study, and subjected to sexing and genotyping by Kompetitive Allele Specific PCR (KASP) and PCR. During the second candling (≈ED18), eggs showing embryonic loss were sampled for tissue collection (embryo retrieval) to verify the *in ovo* sexing and genotyping results using a KASP assay. For samples with discrepancies between allantois and blood/tissue typing, sex determination or genotyping was confirmed by standard endpoint PCR. The remaining 211 eggs served as non-punctured controls to assess hatch rates and were temporarily kept outside the incubator for the same length of time as eggs submitted to sampling. After hatching, blood samples were collected from the 7-day-old chicks that had been subjected to *in ovo* sexing. Blood samples were collected from the Vena metatarsalis plantaris superficialis medialis using a lancet and collected on filter paper (Fig. S3), and subjected to sexing and genotyping by KASP after DNA extraction.

### Phase II Study

450 brown eggs collected from 63-week-old hens, and 450 white eggs collected from 42-week-old hens, sourced from commercial layer flocks, were collected. All 900 eggs were then incubated under standard conditions in an auto-rotating incubator (EMKA Hatchery Equipment B.V., Kuurne, Belgium) at Lohmann Breeders GmbH. White eggs were stored for six days and brown eggs were stored for nine days before incubation was initiated.

At ED4 to ED7, approximately 50 µl of fluid was collected from at least 60 eggs per line per day, using the procedure described in the Pilot study; exact incubation lengths are listed in Table 3. Samples were subjected to sexing and genotyping by KASP assay and standard endpoint PCR. Eggs were candled at ED4, ED7, and again around ED18, a common practice in poultry husbandry before transferring eggs from the setter to the hatcher. After hatching, chicks were sexed visually (vent sexing) by an experienced chick sexer, a standard procedure in commercial poultry facilities. Non-punctured, untreated eggs obtained from 63- and 42-week-old hens, respectively, were used as hatching controls. The brown layer groups represented a cumulative total from three hatches, while the white layer control groups represented a cumulative total from five hatches.

### Phase III Study

A total of 101 iCaspase9 eggs collected across three independent trials (from 43-46-week-old hens) were incubated for 190 hours under standard conditions in a fully automated digital incubator (Ova-Easy 100 Advance Series II Cabinet Incubator, BRINSEA PRODUCTS LTD, North Somerset, UK). Eggs were collected over 7 days. The eggs were obtained from natural mating between: 1) a heterozygous iCaspase9 rooster and iCaspase9 homozygous and heterozygous hens, and 2) a homozygous iCaspase9 rooster and wild-type L68 hens (New Hampshire descent). The air chamber and embryo location were identified using a LED light egg candler. Sterile, single-use lancing devices (ACCU-CHEK, Safe-T-Pro UNO, Roche AG, Basel, Switzerland) were used to puncture the eggshell membrane as described previously^42^ (Fig. 2C). For eggs with a harder shell, a stainless steel needle (T-Pin) was used to create the initial hole in the shell. The fluid was then extracted using a single-use insulin syringe with needle (Covertus BV, Cuijk, Netherlands; Fig. 2D-E). Approximately 50 µl (up to a maximum of 100 µl) of fluid was collected from all fertilized eggs as described in the Pilot study and subjected to sexing and genotyping by PCR. The workflow is illustrated in Fig. 2. After sample collection, eggs were returned to the incubator until ED11, when survival rates post-puncture were assessed. Embryos were then dissected and tissue samples were frozen at -20°C for subsequent sexing and genotyping by PCR. The experiment was repeated using 141 iCaspase9 fertilized eggs. Eggs were collected and subsequently stored at 15°C with 75% relative humidity for a maximum of three weeks. Allantoic fluid was collected from all 141 eggs by puncturing at ED7. Subsequently, 29 eggs of the desired sex and genotype were incubated to hatch. Eggs were moved to the hatcher (Ova-Easy Advance Series II Hatcher, BRINSEA PRODUCTS LTD, North Somerset, UK) at 18 days and 4 hours and incubated at 37.3°C and 65-70% relative humidity. CAM samples were collected from all eggs in the hatcher. Chick quality was assessed using the Pasgar©Score and Tona Score^53,54^.

### DNA Processing and Quantification

DNA was extracted from *in ovo* or allantoic fluids using the REPLI-g Mini Kit (Qiagen, Hilden, Germany) following the manufacturer’s protocol with slight modifications. Samples were stored at -20°C, then thawed on ice and subsequently centrifuged at 13,000 rpm for 1 minute. The supernatant was removed, ensuring the non-visible pellet was undisturbed. Representative samples were stained using Hoechst to demonstrate the presence of cells and/or DNA within the obtained fluid. For Hoechst staining, we resuspended the pellet in 10 µl of Hoechst 33342 solution (5 µg/ml). This was then transferred to a microscope slide, covered with a coverslip, and evaluated using fluorescence microscopy (Zeiss Axioscope 5 with Colibri 5 LED light and Axiocam 202 mono), employing a multi-band pass filter (FS 90 HE LED; Carl Zeiss GmbH, Oberkochen, Germany) and 385/30 nm BP excitation for visualization. For whole genome amplification (WGA), the pellet was resuspended in 3 µl of phosphate buffered saline (PBS) and stored on ice. Incubation with the REPLI-g master mix was carried out for 16 hours using a heated lid thermocycler (Eppendorf Mastercycler) with the following cycling parameters: 16 cycles of 60 minutes at 30°C, 3 minutes at 65°C, and a final hold at 4°C. The heated lid temperature was set to 70°C. For tissue samples, the DNeasy Blood & Tissue Kit (Qiagen) was used. Blood samples were collected on medium filtration capacity filter paper cards^55^ (FTA® card, Schleicher & Schuell GmbH, Whatman, Inc., Maidstone, UK). DNA was then extracted using a pronase digestion protocol as described^56^. CAM and other embryonic tissue were digested by overnight incubation with proteinase K (10 mg/ml) in lysis buffer (100 mM Tris-HCl, 100 mM NaCl, 100 mM EDTA, 1% SDS) at 56°C. After removal of precipitated proteins using saturated NaCl solution and centrifugation, DNA was precipitated with 100% ethanol. The resulting DNA pellet was washed twice with 70% ethanol, air-dried, and dissolved in 30 µl H_2_O. DNA concentration was measured using a FLUOstar Omega (BMG Labtech, Ortenberg, Germany) or Qubit 4 fluorometer (Thermo Fisher Scientific, Waltham, USA) respectively. The concentration of DNA amplified with the REPLI-g Mini Kit was determined using fluorescent dye-based quantification assays, either the Quant-iT PicoGreen dsDNA Assay Kit (Thermo Fisher Scientific, Waltham, USA) or the Qubit 1X dsDNA HS Assay Kit (Invitrogen, Waltham, USA). DNA from WGA samples was visualized on a 0.8% agarose gel (Agarose NEEO ultra-quality, Carl Roth GmbH + Co. KG, Karlsruhe, Germany) in 1x TBE Buffer. Fluorescent staining of nucleic acids in agarose gel was done with ROTI GelStain (Carl Roth GmbH + Co. KG, Karlsruhe, Germany). Signals were visualized using a Fusion SL4-3500.WL (Vilber, Eberhardzell, Germany) UV transilluminator and FusionCapt V15.18 software with a 1.5-second exposure time.

REPLI-g DNA that failed to give a PCR result were purified by ethanol precipitation and the PCR reaction was repeated.

### Sexing and Genotyping by PCR

#### Sexing

Sex determination was performed using a multiplex PCR method modified from Fridolfsson and Ellegren (1999) and a KASP assay (LGC Genomics GmbH, Germany, Berlin), which is a fluorescence resonant energy transfer (FRET) cassette based assay. The KASP assay utilizes allele-specific forward primer to target an A/G difference in exon 17 of the conserved chromodomain helicase DNA binding protein 1 (*CHD1*) gene on the W- and Z-chromosomes (*CHD-Z*/*CHD-W*; Table S1) as described^57^. The multiplex PCR assay exploits an intronic length polymorphism between the CHD-Z and CHD-W^58^. 10 ng of DNA was added to the PCR master mix for each sample.

#### Genotyping

Samples were genotyped for the blue egg (Pilot study and Phase I study) and the iCaspase9-GFP-*DAZL*^23^ locus (Phase III study).

The causal mutation of dominant blue egg shell color in the Araucana breed is a 4.2 kb retroviral insertion (EAV-HP) on chromosome 1 upstream of *SLCO1B3* at 65.22 Mb, which was detected by PCR according to Wragg et al. (2013)^59^ and previously described by our group^60^. Samples from Phase Study III were genotyped for the iCaspase9-GFP integration site in the DAZL locus to distinguish between wildtype, heterozygous, and homozygous carrier animals as described by Ballantyne et al. (2021). PCR amplification was performed using Promega GoTaq Polymerase (Promega, Madison, USA) under the following conditions: initial 95 °C for 2 min, 94 °C for 30 s, 64 °C for 30 s, 72 °C for 30 s for 35 cycles and a final extension of 72 °C for 5 min. Reaction products were resolved using a 1.5 % ultrapure agarose (Invitrogen, Waltham, USA) gel electrophoresis run at 80 V for 45 min in 1× TBE-buffer and visualized using a UV transilluminator. 10 ng of DNA was added to the master mix for each sample.

Each KASP reaction contained about 20-50 ng template DNA, KASP V 4.0 2x Master mix standard ROX (LCG Genomics, Berlin, Germany) and KASP-by-Design assay mix (LGC Genomics). Standard KASP thermal cycling conditions, as described in LGC protocols, were performed using an Eppendorf Mastercycler (Eppendorf, Hamburg, Germany). KASP PCR cycling (blue eggshell) consisted of an initial denaturation step at 94°C for 15 mins, followed by 10 touchdown cycles (94°C for 20 s, 61-55°C for 60 s, with a 0.6°C decrease per cycle). This was followed by 26 additional cycles at 94°C for 20 s and 55°C for 60 seconds. Following amplification, microplates were analyzed with FLUOstar Omega (BMG Labtech, Ortenberg, Germany). Fluorescence signals were detected at excitation/emission wavelengths of 485/520 nm for FAM-labelled-FRET-cassettes, 530/560 nm for HEX-labelled-FRET-cassettes, and 584/620 nm for ROX standard.

#### Data availability statement

Additional data or detailed methods can be obtained from the corresponding author upon reasonable request.

#### Statistical analysis

Statistical analyses were performed with the R statistical software (v.4.0, R Core Team 2023) using the Pearson’s Chi-squared test with Yates’ continuity correction and Fisher’s exact test. Data were considered to be significantly different when P < 0.05.

## Results

### Allantois formation in shell-less culture system

To visualize embryonic development and assess the size of the amnion, allantois and CAM at the time of sample collection, an *ex ovo* culture system adapted from existing methodologies^50,51^ was used (Fig. S4). At 65 hours of incubation (ED2; Fig.S4A), the amnion, which will ultimately enclose the embryonic body, was not yet fully closed. The amniotic fold remained visible, positioned approximately at the level of the posterior vitelline artery (Fig. S4B). By ED3 (72 h; Fig. S4C), the amnion appeared as a small vesicle, originating as an evagination from the endodermal hindgut (Fig. S4D). At ED4 (96 h), the allantois presented as a vesicle, roughly equivalent in size to the embryo’s midbrain (Fig. S4E). At ED4 (116 h; Fig. S4F-H), the allantois was observed to have a mean diameter of 1.67 cm (SD=0.21 cm), while the embryos themselves had a mean size of 1.47 cm (SD=0.09 cm; HH stage 25-26, n=10). At ED6 (144 h; Fig. S4I-K), the size of the allantois increased to 3.1 cm (SD=0.39 cm) with a mean embryo size of 2.4 cm (SD=0.48 cm; HH stage 28-29, n=10). By ED7 (168 h; Fig. S4L-N), the allantois reached a diameter of 4.9 cm (SD=0.57 cm; HH stage 30-32, n=10). This allantoic sac further expanded to 6.6 cm (SD=0.64 cm; HH stage 33-34, n=10) at ED8 (192 h; Fig.S4O-Q), by which time it largely covered the yolk in most of the embryos. At ED9 (240 h; Fig. S4R-T), the allantois entirely covered the yolk and extended beyond it (HH stage 35-36, n=10), as also illustrated in Fig. 3. By ED10 (264 h; Fig.S4U) and ED11 (280 h; Fig. S4V), the allantois contained a greater volume of fluid in all inspected embryos (HH stage 37-38, n=10). In all photographs taken, the embryo is clearly visible within the amniotic cavity, which enlarged concurrently with the growth of the embryo (Fig. S4).

**Fig. 3.**
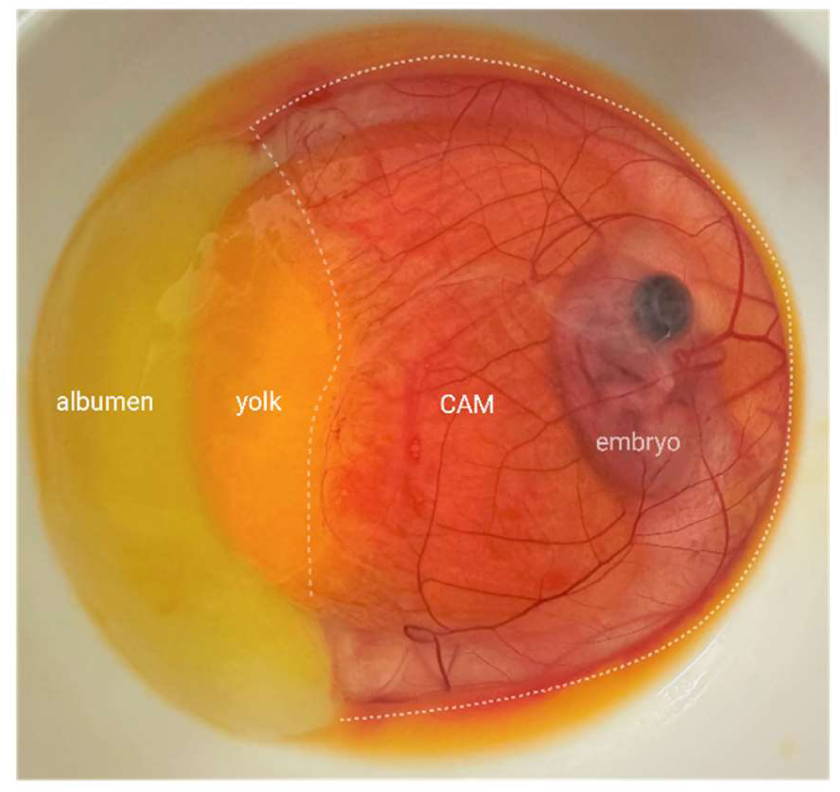
Chicken Embryo and CAM Development at ED9. The highly vascularized chorioallantoic membrane (CAM) encloses the embryo within the amnion and partially covers the yolk sac from all sides at ED9 (indicated by white dashed lines). The CAM is still intact in the depicted image; however, it typically tears when opening the egg shell due to its close connection to the inner shell membrane.

Given the findings described above (Fig. S4E-N), from here on samples collected between ED4 and ED7 of incubation will be referred to as *in ovo* fluid, while samples collected after ED7 of incubation will be referred to as allantoic fluid.

### Pilot study

Across all incubation days (ED4, ED6, ED8, and ED10), 17.5% of the initial shell punctures were unsuccessful, requiring more than one attempt to collect *in ovo* or allantoic fluids. However, sample collection was successful for all eggs. A closer examination of the fluids collected at ED4 and ED7 revealed the presence of very few stainable cells in the sediment (Fig. S5). Direct use of allantoic fluid, simple cell pellet boiling, and standard DNA extraction kits failed to provide measurable DNA yields or consistent PCR results.

WGA yielded high-quality DNA of high molecular weight (Fig. S6). Mean DNA concentration at ED4 was 65.9 ng/µl and remained stable between 72.1 and 75.9 ng/µl from ED8 to ED10 (Table S2). Three samples from early ED4 (96h) showed no detectable DNA on agarose gel. Multiplex PCR-based sexing and genotyping and subsequent gel electrophoresis were successful from the earliest sampling point (ED4) onwards. However, the success rate was lowest at ED4, with 80% for sexing and 67% for genotyping. At ED6, ED8, and ED10 success rates reached 97-100% for sexing and 93%-100% for genotyping, respectively (Table S2). KASP-based sexing and genotyping were successful for all allantoic fluid samples collected at ED8 and ED10 (Table S2). The lowest success rates were observed at ED4 (70% for sexing and 73% for genotyping). At ED6, success rates increased to 97% for sexing and 93% for genotyping.

Sexing and genotyping were successful for all embryonic tissue control samples. Discordant results were observed for sexing and genotyping in two fluid samples from ED6 when compared to the results from the control samples. For ED10 control samples, sex was additionally verified by macroscopic examination of the developing gonads (Fig. S2A-D), confirming the sex previously determined *in ovo*. The results of the allelic discrimination assay, which was used for sexing and genotyping the blue egg allele, are exemplified by the cluster plot in Fig. 4.

**Fig. 4.**
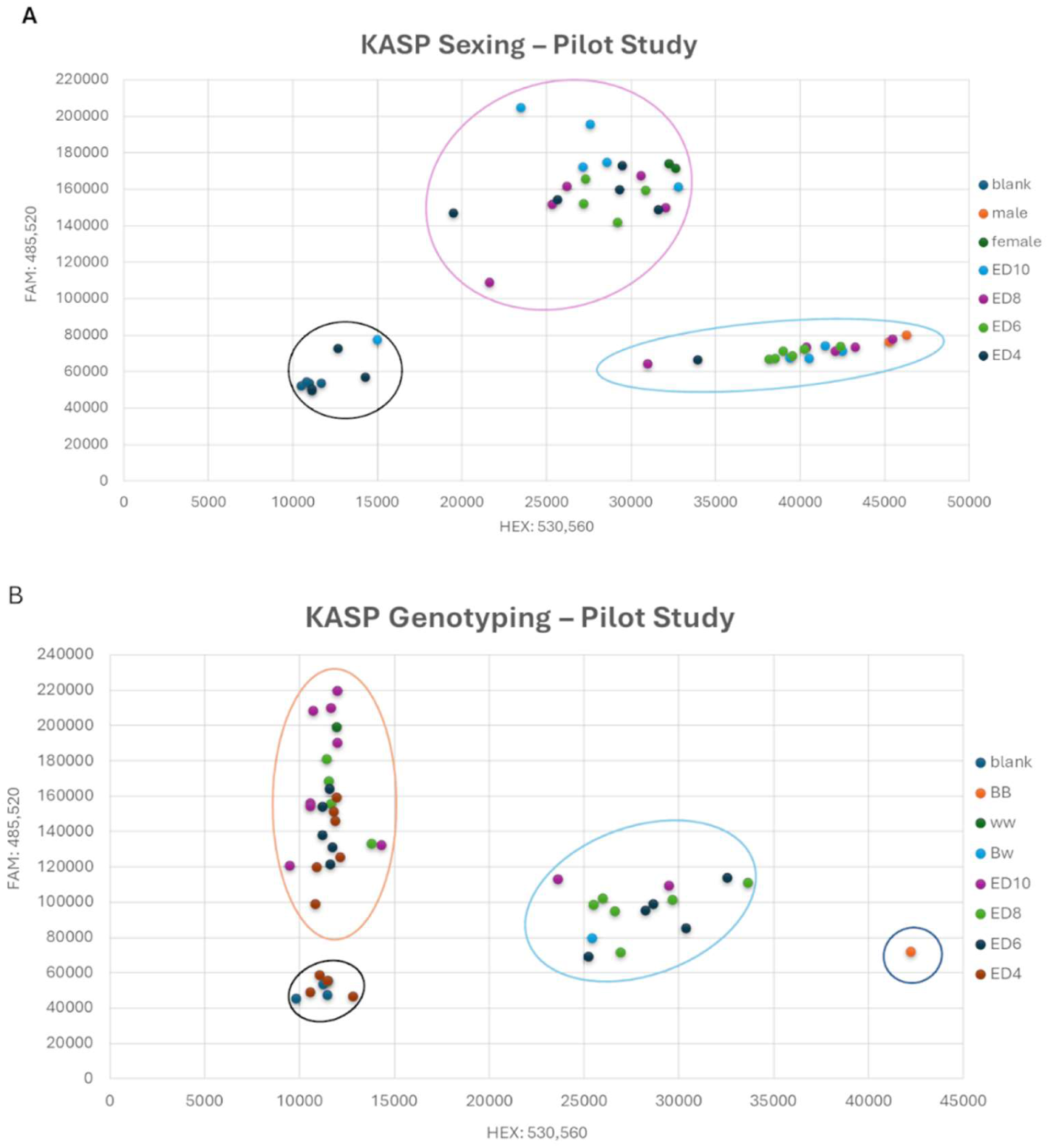
Fluorescence-based Allelic Discrimination using DNA from *in ovo* or Allantoic Fluid. Representative cluster plots for the Kompetitive allele specific PCR (KASP) sexing and genotyping assay (Pilot study) using whole-genome amplified (WGA) DNA extracted from *in ovo* or allantoic fluid. **(A) Sexing cluster plot**: The female-specific forward primer signal (FAM) is plotted against the male-specific forward primer signal (HEX). Distinct clusters represent females (circled in pink) and males (circled in blue), including samples collected at ED4, ED6, ED8 and ED10. The plot also includes reference samples for validation, as well as a no-template control (NTC) cluster (circled in black). The clear separation of these clusters demonstrates the successful differentiation between sexes. A few samples failed to amplify (clustering with NTC) and were subsequently re-analyzed. **(B) Genotyping cluster plot**: The plot displays three distinct clusters: BB (homozygous blue, circled in dark blue), Bw (heterozygous blue, circled in light blue) and ww (wild-type, circled in orange). No-template controls are also shown and circled in black. Reference samples (blood-derived) are labeled as BB, Bw, and ww, while the test samples from ED4, ED6, ED8, and Ed10 are labeled accordingly.

### Phase I study

Of the 480 eggs in the study, 52 eggs were excluded prior to allantoic fluid collection (39 were unfertilized and 13 embryos were determined not-viable at the first candling at ED7). Of the remaining 428 eggs, 217 were sampled at ED7 (184 hours), while 211 served as control to assess the impact of egg puncture on hatchability. Twenty-eight percent of the initial shell punctures were unsuccessful, requiring more than one attempt to collect allantoic fluid. The eggs were candled at ED18 to count and remove those with dead-in-shell embryos, and non-hatched chicks were also examined to determine the day of death. Two embryos in each group (control and treated) were identified as early embryonic losses (≤ED 6), having been overlooked at the first candling at ED7 (Table 1). Four embryos in the control group and eleven in the sampled group were classified as mid-embryonic death (ED7-ED12), occurring shortly after sample collection (Table 1). Most dead embryos were categorized as late embryonic deaths (ED13-ED20), with the highest number of deaths occurring between ED18-ED20. Late embryonic deaths totaled 24 in the sampled group and 13 in the control group. The mid and late embryonic mortality was higher in the sampled group (16%) compared to the control group (8%). In the control group, 90% of eggs yielded healthy hatchlings, compared to 82% in the sampled group (Table 1). The percentage of live chicks in the treated group (86%) was significantly lower than in the control group (90.9%; χ²(1)=5.7992, p=0.01603).

**Table 1.** Hatching Results for Phase I study.

| Group | Total number of eggs | Number of unfertilized eggs or early losses (<ED7) | Number of mid embryonic deaths (ED7-12) | Number of late embryonic deaths (ED13-20) | Number of unhatched eggs (ED7-20) | Number of hatched chicks | Number of chicks lost post-hatch | Number of healthy hatchlings |
| --- | --- | --- | --- | --- | --- | --- | --- | --- |
| Samples | 217 | 2 | 11 | 24 | 35 (16%) | 180 | 1 (0.5%) | 179 (82%) |
| Controls | 211 | 2 | 4 | 13 | 17 (8%) | 192 | 2 (1%) | 190 (90%) |
| Total | 428 | 4 | 15 | 37 | 52 (12%) | 372 | 3 (0.8%) | 369 (86%) |

Further analysis of embryonic death revealed that the difference in mid embryonic death (Table 1; punctured: 5.1%, n=11/215 vs. non-punctured: 1.9%, n=4/209) was not statistically significant (Fisher’s Exact Test, p=0.113). Similarly, the difference in late embryonic death (Table 1; punctured: 11.2%, n=24/215 vs. non-punctured: 6.2%, n=13/209) also did not reach statistical significance (Chi-square test, χ^2^(1)=2.66, p=0.103).

215 samples were used to validate the accuracy of sexing and genotyping from allantoic fluid DNA subjected to both endpoint and multiplex PCRs with gel electrophoresis and fluorescence-based KASP assays. Results were compared with KASP assay outcomes from DNA extracted from blood of corresponding hatched chicks and embryonic tissue for non-viable embryos. While all KASP assays yielded results, 17.6% of the allantoic fluid DNA samples required repeat KASP sexing due to unclear clustering, and 5.11% required repeat KASP genotyping. On the other hand, standard PCR achieved success rates of 99.5% for sexing and 99% for genotyping in terms of concordance (Table 2). Comparing KASP sexing results from allantoic fluid to those obtained for tissue/blood samples revealed high accuracy (99.5), with only two samples being mismatches. For KASP genotyping, the accuracy was 95%, with ten samples displaying a mismatch. Concordance between standard PCR genotyping of allantoic fluid and KASP genotyping of tissue/blood samples was 96%, with seven mismatches. Standard PCR sexing of allantoic fluid displayed 99% concordance with KASP sexing of tissue/blood controls, with two mismatches. Finally, direct comparison between standard PCR and KASP from allantoic fluid yielded 99% accuracy for sexing (two mismatches) and 98% accuracy for genotyping (four mismatches; Table 2).

**Table 2.** Sexing and EAV-HP-Insertion Genotyping Success and Accuracy Rates (Phase I study – ED7)

| Sample size total/ relative | Number of successful Sexing (PCR) | Number of successful Sexing (KASP) | Number of Concordance Sexing KASP (AF) vs. PCR (AF) | Number of Concordance Sexing AF (KASP) vs. control (KASP) | Number of Concordance Sexing AF (PCR) vs. control (KASP) | Number of Successful Genotyping (PCR) | Number of Successful Genotyping (KASP) | Number of Concordance EAV-HP Genotyping KASP (AF) vs. PCR (AF) | Number of Concordance EAV-HP Genotyping AF (KASP) vs. control (KASP) | Number of Concordance EAV-HP Genotyping AF (PCR) vs. control (KASP) |
| --- | --- | --- | --- | --- | --- | --- | --- | --- | --- | --- |
| 215 | 214 (99.5%) | 215 (100%) | 213 (99%) | 213 (99%) | 213 (99%) | 214 (99%) | 215 (100%) | 211 (98%) | 205 (95%) | 208 (96%) |

AF= Allantoic fluid. control = blood from hatched chicks or tissue from non-hatched chicks but punctured eggs.

### Phase II study

Most egg fluid samples obtained at ED7 were clear, while samples collected from ED4 to ED6 were frequently cloudy and yellowish (Fig. S7). Only one sample appeared to be bloody (Table 3). Success rates of fluid collection varied depending on the day of incubation. Remarkably, at ED6, up to 25% of the eggs were excluded because fluid could not be successfully extracted despite numerous puncture attempts. However, at least 60 samples from this trial were successfully punctured.

**Table 3.**
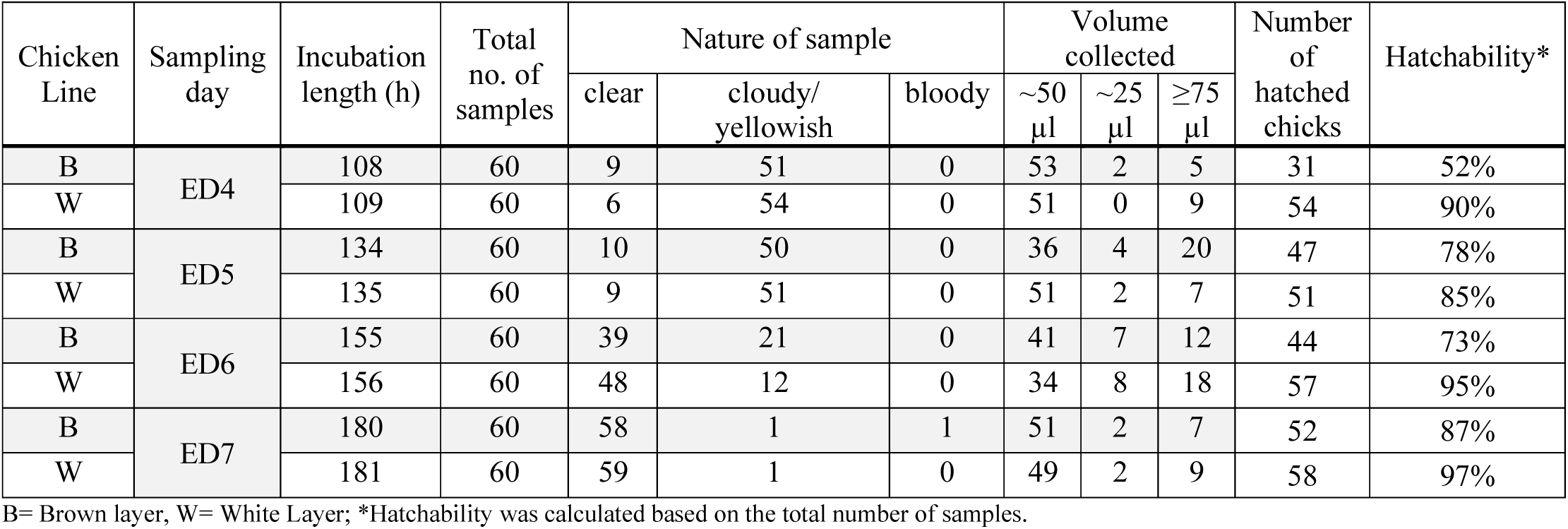
Sample Data for Phase II Study (Chick Sexing ED4-ED7)

Following sample collection at ED4, brown eggs exhibited a hatch rate of 52%, while white eggs showed a higher rate of 90% (Table 3). After sample collection at ED5, the hatch rates for brown eggs increased to 79% and white eggs reached 85%. Sample collection at ED6 led to a slight decrease in the hatch rate for brown eggs (73%), while the hatch rate for white eggs remained high (95%). Finally, following sample collection at ED7, the hatch rates were 87% for brown eggs and 97% for white eggs, displaying the highest rates observed in the study (Table 3).

The overall hatch rate for *in ovo* sexed brown eggs was 73%; of those 53% were males and 47% were females (Table S3). White eggs sexed *in ovo* achieved a 92% hatch rate (50% males and 50% females). Unfertilized eggs were excluded prior to sampling. In the control hatch, incubated simultaneously to the *in ovo* sexed group, the unfertilized egg rate was 26% for brown eggs and 6% for white eggs. Regrettably, a subset of eggs from the flock control group was subjected to *in ovo* sexing technique evaluation but not excluded from the hatch. The hatch rate of brown eggs from the control flock was 63% (52% males and 48% females), while the hatch rate for white control eggs was 92% (54% males and 46% females). Routine hatchery (utilizing hens of identical age) not subjected to *in ovo* sexing demonstrated hatch rates of 37-41% for brown layers (three separate hatches) and 44-47% for white layers (five separate hatches; Table S2). These groups exhibited non-fertilized egg rates of 12-16% for the brown chicken line and 2-7% for the white line.

KASP sexing demonstrated an overall success rate of 92% across all incubation days (362 out of 394 samples; Table 4). Success rates varied by egg type and incubation day: at ED4, KASP achieved 77% success for brown eggs and 87% for white eggs. By ED5, success rates increased to 91% for brown eggs and 86% for white eggs. ED6 yielded further gains, with 98% success for brown eggs and 93% for white eggs. At ED7, the success rate reached 96% for brown eggs and 100% for white eggs. While the overall concordance between KASP results and vent sexing post-hatch was 98% across all incubation days, concordance was slightly lower for both brown and white chicks at ED4 and ED7 (92-96%; Table 4).

**Table 4.** Success Rates of Sex Determination using KASP Assay on *In Ovo* Fluid or Allantoic Fluid Samples (Phase II study)

| Chicken Line | Sampling Day | Incubation length (h) | Number of Hatched & Sexed Chicks | Number of Successful KASP Sexing | Concordance KASP & Sexer (N) | Concordance KASP & Sexer |
| --- | --- | --- | --- | --- | --- | --- |
| B | ED4 | 108 | 31 | 24 (77%) | 23 | 96% |
| W |  | 109 | 54 | 47 (87%) | 45 | 96% |
| B | ED5 | 134 | 47 | 43 (91%) | 43 | 100% |
| W |  | 135 | 51 | 44 (86%) | 44 | 100% |
| B | ED6 | 155 | 44 | 43 (98%) | 43 | 100% |
| W |  | 156 | 57 | 53 (93%) | 53 | 100% |
| B | ED7 | 180 | 52 | 50 (96%) | 46 | 92% |
| W |  | 181 | 58 | 58 (100%) | 56 | 96% |
| Total | ED4-7 | 108-181 | 394 | 362 (92%) | 353 | 98%** |

B= Brown layer, W=White layer, KASP: Kompetitive allele-specific PCR; Sexer: Sex determination by visual inspection; KASP assay sexing was performed only on samples from live, sexed chicks. \*\*Mean %.

### Phase III study

Across all experimental groups, 101 iCaspase9 eggs were punctured at ED7, 26% of which were punctured at least twice to aspirate fluid. The aspirated fluid was bloody in four samples, clear in 65 samples, and yellowish in 31 samples. The survival rate at ED11 was 94% in the first replicate, 88% in the second replicate, and 94% in the third replicate (Table S4). Two embryos died during the first replicate: one at ED8 and another with a bloody amnion at ED9. The second replicate recorded four embryonic deaths: two occurred before ED9 (prior to puncturing). Additionally, one death occurred at ED11 and another at ED9, both exhibiting blood in the amniotic cavity. During the third replicate two embryos died at ED9. Among the untreated control eggs incubated simultaneously, four embryonic deaths occurred, all before ED8. Sexing and genotyping results for the iCaspase9-GFP transgene were obtained for all aspirated samples. However, one result each from the sexing and genotyping deviated from the tissue analysis (Table S4). The male to female embryo ratio, approximately 50:50, along with the identified genotypes for each trial, are presented in Table S4. As expected, based on the crossing scheme, most embryos were identified as heterozygous iCaspase9 carriers.

The second experimental phase, which included incubation to hatch, involved a total of 177 iCaspase9 eggs. Of these, 141 were confirmed as fertilized by ED7, three died prior to ED7, and the remainder (n=33) were unfertilized. Of the 141 allantoic fluid samples collected at ED7 and initially tested for sexing and genotyping by PCR, 19 did not yield results. Following purification, all but three of the samples produced results (98%). Three mismatches were observed between the allantois and control tissue samples, two for genotyping and one for sexing (Table S5).

29 iCaspase9 eggs of the desired sex and genotype were selected for full-term incubation. All selected eggs were confirmed viable by candling at ED18, and subsequently transferred to the hatcher. A total of 25 chicks successfully hatched. One chick died two days post-hatch due to poor development and open navel at hatch. The remaining chicks were healthy and developed normally (Fig. S8). Specifically, the 29 eggs chosen for hatching comprised: one heterozygous and five homozygous iCaspase9 males, and seven wild-type, 15 heterozygous and one homozygous iCaspase9 females. Of these, one heterozygous and three homozygous males, and seven wild-type and 13 heterozygous females successfully hatched and survived (24 chicks in total: 4 males and 20 females). One heterozygous female chick died in week four (unknown cause). Examination of the five non-hatched chicks showed that they were correctly positioned and developed within the egg but had not initiated internal pipping into the air chamber. Sex and genotypes were confirmed by PCR analysis of allantoic fluid and corresponding post-hatch CAM tissue samples (Fig. S9). Egg weight loss at ED18 averaged 12% (SD=2%) compared to initial egg weight. The mean hatch weight for chicks of all sexes and genotypes was 41.7g (SD=3.3g). Chicks reached a mean body weight of 253.9g (SD=31.4g) by four weeks and 581.6g (SD=61.9g) by six weeks of age. Their growth performance matched expectations when compared to the parental iCaspase9 chicks (Fig. S10).

## Discussion

In this study, we successfully developed a reliable and user-friendly method for *in ovo* sexing and genotyping of different chicken lines used for research purposes. Our results confirmed the feasibility of *in ovo* sexing and genotyping from ED4 (96h of incubation) to ED10 (240h of incubation), and identified an optimal window for allantoic fluid collection which begins at ED7 for maximal reliability, minimal embryonic impact and sufficient hatchability. The choice of ED7 balances this optimal performance window with the requirement to provide sufficient time for DNA analysis before the assumed onset of first signs of embryonic nociception (ED13)^10–12^.

Our shell-less culture system demonstrated the rapid development and expansion of the allantois, a finding consistent with observations in dynamic 3D magnetic resonance imaging (MRI) studies^61^. In early stages (ED2-ED6), the allantois remains a small vesicle, making targeted puncture through the eggshell with a short cannula exceptionally challenging. This is reflected by our observation that fluid collected prior to ED7 was predominantly yellowish rather than clear. We propose that fluid aspirated prior to extended allantois and CAM formation (<ED7) originates from the sub-embryonic fluid (SEF), located within the yolk sac beneath the embryo^62^. While DNA extraction and subsequent sex/genotype determination was feasible with this early fluid, success rates were lower between 96 and 144 hours of incubation (ED4-ED6) compared to later stages.

Our observations are supported by the volume dynamics described in the literature. The total SEF volume in the chicken egg reaches a maximum of 13 ml at ED6 and is subsequently reabsorbed until ED15^62,63^, aligning with our data that ED6/ED7 appears to be a turning point. Empirically, this transition is reflected in the high rate of challenging aspirations, which in our study required multiple attempts in 18-28% of eggs across all of our studies. The difficulty in aspirating fluid peaked at ED6 (155-156h), resulting in 25% of eggs being rejected at this time point during the Phase II study.

We suggest that this critical time point reflects the transition from SEF to allantoic fluid, a stage where the developing allantoic sac contains less targetable fluid. This aligns with external reports that sample collection at early stages results in the highest rate of embryo death, likely due to the increased difficulty of the procedure^42^.

Given the crucial role of SEF in early embryonic viability, including potential contributions to acid-base regulation and tissue hydration^64^, our data suggest that the risk of compromising embryonic development through fluid extraction decreases as incubation progresses. Higher hatch rates in eggs punctured later in development compared to those punctured earlier (Phase II study) are an indicator of this reduced risk.

Previous studies have shown that allantoic fluid volume sharply increases from ED7 onwards, following a parabolic-shaped profile that peaks at ED12^61,62^. We hypothesize that this augmented fluid volume acts as a protective buffer, minimizing the potential for damage to embryonic development and extraembryonic membrane expansion. We visually confirmed that, at ED7, the fluid was exclusively present in the allantoic sac, not in the amnion. This accumulation is consistent with the allantois serving as a repository for embryonic kidney excretions, which begin around ED5^62^. Like amniotic fluid (6 ml), allantoic fluid reaches its maximum water content (≈8-14 ml) at ED12/13^61–63^. Concurrently, the CAM serves as the embryo’s primary respiratory surface during the latter half of incubation^62^. Avoiding disruption of the developing CAM through puncture is thus critical.

Despite initial findings suggesting a negative impact of the puncture procedure on hatch rates, the larger Phase II study (conducted at the optimal ED7 timepoint) found no evidence that hatchability was impacted by puncture. These conflicting results can most likely be attributed to the limited statistical power to detect the specific effects of early and late mortality phases individually in the Phase I study, even though they contributed to the significant overall effect. We hypothesize that mid embryonic death observed in the Phase I study is specifically associated with the puncture-induced trauma, a conclusion supported by the Phase III study, which documented embryonic death as late as ED8, presenting with blood-filled amnion. Late embryonic death typically occurs around ED19 due to factors such as malposition, membrane entrapment^65^ and failure to transition from allantois to pulmonary respiration^66^. Since a detailed investigation of these factors was beyond the scope of this study, the potential contribution of our puncture procedure to late embryonic mortality cannot be ruled out. Although the Phase II and Phase III studies showed limited impact on hatchability when our method was performed at ED7, ongoing concerns from breeding companies regarding potential puncture-induced undesirable side effects (e.g. disturbed embryo development or contamination) have already led to the development of non-invasive *in ovo* sexing methods, as previously reviewed elsewhere^14^. High-quality DNA obtained from *in ovo* fluids by WGA enabled the implementation of various assays, including fluorescence-based sexing and genotyping as well as multiplex PCR followed by gel electrophoresis. Our accuracy rates, ranging from 92-100% across all incubation days, are comparable to other established methods. This includes robotic, large-scale *in ovo* sexing system like that used used by PLANTegg GmbH^41^ (≈ 99.5% accuracy at ED9) and published qPCR assays utilizing allantoic fluid DNA (95-100%)^42^. Crucially, the Pilot and Phase II studies clearly demonstrated that PCR success rates improved with increasing embryonic age, likely due to higher cell/DNA content in the samples. While our method achieved high accuracy, minor discrepancies between sample and control results were observed (e.g., allantoic fluid samples being genotyped as male when tissue controls were female). These are primarily attributed to inherent limitations such as human handling error during fluid collection or assay variability potentially caused by PCR inhibitors derived from the collected fluid.

Our findings confirm that performing the invasive puncture and fluid extraction at ED7 has limited impact on embryo survival and hatchability, consistent with prior research^37,38,42^. Our four independent studies, which utilized various breeds, multiple laboratories with varied equipment, demonstrate the method’s feasibility under diverse conditions. These results confirm that *in ovo* sexing and genotyping can be performed by smaller laboratories equipped with a PCR cycler or fluorometer, suggesting that this method can be rapidly adopted after brief training.

## Conclusion

Reducing the number of surplus chicks through early sexing and genotyping saves labor and resources. These *in ovo* techniques also have the potential to be applied to mammalian models, for example by puncturing blastocysts to obtain blastocoel fluid for sex and genotype determination in cattle and pigs^67^. A critical need exists to re-evaluate the ethical framework and enhance animal welfare standards for farm animals utilized in research, such as chickens, pigs, and cattle. By responsible implementation of innovative approaches, we can leverage the chicken as an animal model while upholding ethical standards. It requires enhanced understanding of nociception and analgesic effects in chickens, alongside more objective pain assessment for all avian species. Consequently, there is a strong drive to develop robust anesthesia and analgesia protocols for chickens and their embryos for use during potentially painful procedures^68–73^. The recently introduced ‘Stress Chicken Scale’ by Schlegel et al. (2024) highlights the need for effective pain recognition tools for adult birds, thus demonstrating the relevance of pain management beyond the embryonic stage^74^.

## Limitations of the study

Not all groups included in this study were brought to hatch, precluding an assessment of impact on hatchability. Additionally, follow-up monitoring of adult chickens (e.g. growth curves or laying performance) was not conducted. As sex determination in the Phase II study relied solely on vent sexing, it is not possible to ascertain whether discrepancies between KASP and vent sexing results are due to the sexer’s assessment, or to issues related to PCR or sample handling. Assessments for proper embryo development, including the ratio of embryo or yolk weight to total egg weight, were not part of this study. To enhance the statistical power of the data, future studies would benefit from testing a larger number of flocks and eggs across repeated trials.

## Author contributions

Conceptualization and project design: C.D., S.A., S.W., A.F., D.M. and R.P.; Investigation (egg incubation and candling, *in ovo* extraction of allantoic fluid, DNA extraction, DNA quantification, PCR, staining, microscopy): S.A., C.D., A.F.; Visualization: S.A. and C.D.; Writing: S.A and C.D.; Review and editing: D.M., C.K., S.W., S.A, C.D, R.P. and A.F.; Transfer of the iCaspase9 line^23^ to FLI: D.M. and S.A. All authors read and agreed to the published version of the manuscript.

## Acknowledgements/Funding

We thank Lohmann Breeders GmbH for providing eggs from a backcross population developed under the framework of the EU project IMAGE, which received funding from the European Union’s Horizon 2020 research and innovation programme under grant agreement No 677353. The backcross generation based on Araucana males and White Leghorn females was previously published in a conference abstract^48,49^. D. Meunier and the National Avian Research Facility are funded by the Biotechnology and Biological Sciences Research Council, UK (grant BB/CCG2270/1). We thank Mike McGrew from the Roslin Institute and Royal (Dick) School of Veterinary Studies University of Edinburgh for his support in facilitating the transfer of iCaspase9 eggs^23^ to Germany. The authors are sincerely grateful for the excellent technical assistance of Meike Stünkel and Annett Weigend throughout the study. The authors declare that no competing interests exist.

## Notes

### Competing Interest Statement

The authors have declared no competing interest.

## References

1 Bahr, J. M. in Sourcebook of Models for Biomedical Research (ed P. Michael Conn) 161–167 (Humana Press, 2008).

2 Beacon, T. H. & Davie, J. R. The chicken model organism for epigenomic research. Genome 64, 476–489 (2021). 10.1139/gen-2020-0129

3 Kaplan-Arabaci, O., Dančišinová, Z. & Paulsen, R. E. The Chicken Embryo: An Alternative Animal Model in Development and Disease. Heliyon (2024). 10.2139/ssrn.4724622

4 Burt, D. W. Emergence of the chicken as a model organism: implications for agriculture and biology. Poult Sci 86, 1460–1471 (2007). 10.1093/ps/86.7.1460

5 Stern, C. D. The chick; a great model system becomes even greater. Dev Cell 8, 9–17 (2005). 10.1016/j.devcel.2004.11.018

6 Sarnella, A. et al. The Chicken Embryo: An Old but Promising Model for In Vivo Preclinical Research. Biomedicines 12 (2024). 10.3390/biomedicines12122835

7 Davey, M. G. & Tickle, C. The chicken as a model for embryonic development. Cytogenet. Genome Res 117, 231–239 (2007). 10.1159/000103184

8 Gruber, F. P. & Hartung, T. Alternatives to animal experimentation in basic research. ALTEX (2004).

9 Davey, M. G. a. M., Mike J. and Holmes, Tana. A Scientific Case for Revisiting the Embryonic Chicken Model in Biomedical Research. SSRN (2024). 10.2139/ssrn.5012603

10 Weiss, L. et al. Nocicepton in chicken embryos, Part I: Analysis of cardiovascular responses to a mechanical noxious stimulus. Animals (Basel) 13 (2023). 10.1101/2023.04.14.536899

11 Kollmansperger, S. et al. Nociception in chicken embryos, Part II: Embryonal development of electroencephalic neuronal activity in ovo as a prerequisite for nociception. bioRxiv (2023). 10.1101/2023.04.14.536947

12 Süß, S. C. et al. Nociception in chicken embryos, Part III: Analysis of movements before and after application of a noxious stimulus. Animals 13 (2023). 10.1101/2023.04.20.537674

13 Bruijnis, M. R. N., Blok, V., Stassen, E. N. & Gremmen, H. G. J. Moral “Lock-In” in Responsible Innovation: The Ethical and Social Aspects of Killing Day-Old Chicks and Its Alternatives. Journal of Agricultural and Environmental Ethics 28, 939–960 (2015). 10.1007/s10806-015-9566-7

14 Corion, M., Santos, S., De Ketelaere, B., Spasic, D., Hertog, M. & Lammertyn, J. Trends in in ovo sexing technologies: insights and interpretation from papers and patents. J Anim Sci Biotechnol 14, 102 (2023). 10.1186/s40104-023-00898-1

15 Xu, S., Long, S., Su, Z., Hayat, K., Xie, L. & Pan, J. Egg characteristics assessment as an enabler for in-ovo sexing technology: A review. Biosystems Engineering 249, 41–57 (2025). 10.1016/j.biosystemseng.2024.11.008

16 Di Concetto, A., Morice, O., Corion, M. & Monteiro Belo dos Santos, S. Chick and Duckling Killing: Achieving an EU-Wide Prohibition. (European Institute for Animal Law & Policy, 2023).

17 Code rurale de la pêche maritime; Partie réglemetaire. Journal officiel de la République française, Livre II, Chapitre IV, Section 2, Sous-section 1, R214–217 (2022).

18 Code rurale de la pêche maritime; Partie réglementaire Journal officiel de la République française, Livre II, Chapitre IV, Section 4, Sous-Section 3, R214-278 (modifié par Décret n°2022-2137-Art.2021) (2022).

19. Animals (Scientific Procedures) Act 1986. London Gazette, c. 14 (2013).

20 German Animal Welfare Act (Tierschutzgesetz). BGBl. I, S 1826, §4c (2021).

21 Animal Welfare Regulation Governing Experimental Animals (TierSchVerV). BGBl. I S. 1308, Section 2, §14 (2024).

22 Bacon, L. D., Hunt, H. D. & Cheng, H. H. A review of the development of chicken lines to resolve genes determining resistance to diseases. Poult Sci 79, 1082–1093 (2000). 10.1093/ps/79.8.1082

23 Ballantyne, M. et al. Direct allele introgression into pure chicken breeds using Sire Dam Surrogate (SDS) mating. Nat Commun 12, 659 (2021). 10.1038/s41467-020-20812-x

24 Lengyel, K. et al. Unveiling the critical role of androgen receptor signaling in avian sexual development. Nat Commun 15, 8970 (2024). 10.1038/s41467-024-52989-w

25 Jansen, S. et al. Relationship between Bone Stability and Egg Production in Genetically Divergent Chicken Layer Lines. Animals (Basel) 10 (2020). 10.3390/ani10050850

26 Henderson, L., Okuzaki, Y., Marcelle, C., McGrew, M. J. & Nishijima, K. I. Avian bioresources for developmental biology: Chicken and quail resources in the United Kingdom, France, and Japan. Dev Biol 521, 1–13 (2025). 10.1016/j.ydbio.2025.02.001

27 Milchevskaya, V. et al. Group size planning for breedings of gene-modified mice and other organisms following Mendelian inheritance. Lab Anim (NY) 52, 183–188 (2023). 10.1038/s41684-023-01213-1

28 Buch, T., Davidson, J., Hose, K., Jerchow, B., Nagel-Riedasch, S. & Schenkel, J. Reducing surplus experimental animal generation. Laboratory Animals 56, 305–305 (2022). 10.1177/00236772221096054

29 Wewetzer, H., Wagenknecht, T., Bert, B. & Schonfelder, G. The fate of surplus laboratory animals: Minimizing the production of surplus animals has greatest potential to reduce the number of laboratory animals. EMBO Rep 24, e56551 (2023). 10.15252/embr.202256551

30 Morinha, F. et al. High-resolution melting analysis for bird sexing: a successful approach to molecular sex identification using different biological samples. Mol Ecol Resour 13, 473–483 (2013). 10.1111/1755-0998.12081

31 Cordeiro, C. D. et al. Fast, accurate, and cost-effective poultry sex genotyping using real-time polymerase chain reaction. Front Vet Sci 10, 1196755 (2023). 10.3389/fvets.2023.1196755

32 He, L. et al. Simple, sensitive and robust chicken specific sexing assays, compliant with large scale analysis. PLoS One 14, e0213033 (2019). 10.1371/journal.pone.0213033

33 Rosenthal, N. F., Ellis, H., Shioda, K., Mahoney, C., Coser, K. R. & Shioda, T. High-throughput applicable genomic sex typing of chicken by TaqMan real-time quantitative polymerase chain reaction. Poult Sci 89, 1451–1456 (2010). 10.3382/ps.2010-00638

34 Clinton, M., Haines, L., Belloir, B. & McBride, D. Sexing chick embryos: a rapid and simple protocol. Br Poult Sci 42, 134–138 (2001). 10.1080/713655025

35 Jensen, T., Mace, M. & Durrant, B. Sexing of mid-incubation avian embryos as a management tool for zoological breeding programs. Zoo Biol 31, 694–704 (2012). 10.1002/zoo.20433

36 Hofstadt, M. V. D. et al. Molecular sexing of chick embryos by LAMP and RPA assays: a step toward in ovo egg sexing. PREPRINT (Version 1*) available at Research Square* (2025). 10.21203/rs.3.rs-5772672/v1

37 Weissmann, A., Reitemeier, S., Hahn, A., Gottschalk, J. & Einspanier, A. Sexing domestic chicken before hatch: a new method for in ovo gender identification. Theriogenology 80, 199–205 (2013). 10.1016/j.theriogenology.2013.04.014

38 Weissmann, A. et al. In ovo-gender identification in laying hen hybrids: Effects on hatching and production performance. Europ.Poult.Sci (2014). 10.1399/eps.2014.25

39 Turkyilmaz, M. K., Karagenc, L. & Fidan, E. Sexing of newly-hatched chicks using DNA isolated from chorio-allantoic membrane samples by polymerase chain reaction in Denizli chicken. Br Poult Sci 51, 525–529 (2010). 10.1080/00071668.2010.502521

40 Seleggt. <https://www.seleggt.com/> (Accessed October 28, 2025).

41 PLANTegg. <https://www.plantegg.de/> (Accessed October 28, 2025).

42 Monteiro Belo Santos, S., Corion, M., De Ketelaere, B., Lammertyn, J. & Spasic, D. Allantoic Fluid-Based qPCR for Early Onset In Ovo Sexing. J Agric Food Chem (2024). 10.1021/acs.jafc.3c09418

43 Respeggt. <https://www.respeggt.com> (Accessed October 28, 2025).

44 Woodcock, M. E. et al. Reviving rare chicken breeds using genetically engineered sterility in surrogate host birds. Proc. Natl. Acad. Sci. U. S. A. 116, 20930–20937 (2019). 10.1073/pnas.1906316116

45 Panda, S. K. & McGrew, M. J. Genome editing of avian species: implications for animal use and welfare. Lab Anim 56, 50–59 (2022). 10.1177/0023677221998400

46 Percie du Sert, N., et al. Reporting animal research: Explanation and elaboration for the ARRIVE guidelines 2.0. PLoS Biol 18, e3000411 (2020). 10.1371/journal.pbio.3000411

47 Percie du Sert, N., et al. The ARRIVE guidelines 2.0: Updated guidelines for reporting animal research. PLoS Biol 18, e3000410 (2020). 10.1371/journal.pbio.3000410

48 Dierks C., N. T. H., H. Simianer, D. Cavero, R. Preisinger, S. Weigend. in 70th Annual Meeting of the European Federation of Animal Science, 26–30 August 2019. (ed Scientific Committee).

49 Dierks C. , N. T. H., H. Simianer, B. Andersson, D. Cavero, R. Preisinger, S. Weigend. in XI European Symposium on Poultry Genetics, 23–25 October 2019. (ed S. Weigend P. Trefil).

50 Mangir, N., Dikici, S., Claeyssens, F. & MacNeil, S. Using ex Ovo Chick Chorioallantoic Membrane (CAM) Assay To Evaluate the Biocompatibility and Angiogenic Response to Biomaterials. ACS Biomater Sci Eng 5, 3190–3200 (2019). 10.1021/acsbiomaterials.9b00172

51 Cloney, K. & Franz-Odendaal, T. A. Optimized ex-ovo culturing of chick embryos to advanced stages of development. J Vis Exp, 52129 (2015). 10.3791/52129

52 Hamburger & Hamilton. A series of normal stages in the development of the chick embryo. J Morphol 88(1): 49-92. (1951).

53 Tona, K. et al. Effects of egg storage time on spread of hatch, chick quality, and chick juvenile growth. Poult Sci 82, 736–741 (2003). 10.1093/ps/82.5.736

54 Boerjan, M. L. in Avian and Poultry Biology Reviews. 4 edn 237–238.

55 Suriyaphol, G., Kunnasut, N., Sirisawadi, S., Wanasawaeng, W. & Dhitavat, S. Evaluation of dried blood spot collection paper blotters for avian sexing by direct PCR. Br. Poult. Sci. 55, 321–328 (2014). 10.1080/00071668.2014.925087

56 Bailes, S. M., Devers, J. J., Kirby, J. D. & Rhoads, D. D. An Inexpensive, Simple Protocol for DNA Isolation from Blood for High-Throughput Genotyping by Polymerase Chain Reaction or Restriction Endonuclease Digestion. Poultry Science 86, 102–106 (2007). 10.1093/ps/86.1.102

57 Dierks, C., Altgilbers, S., Weigend, A., Preisinger, R. & Weigend, S. Sexing assay for chickens and other birds for large-scale application based on a conserved sequence variant in CHD1 genes on W and Z chromosomes. Anim Genet 53, 235–237 (2022). 10.1111/age.13176

58 Fridolfsson, A. K. & Ellegren, H. A simple and universal method for molecular sexing of non-ratite birds. J. Avian Biol. 30, 116–121 (1999).

59 Wragg, D. et al. Endogenous retrovirus EAV-HP linked to blue egg phenotype in Mapuche fowl. PLoS One 8, e71393 (2013). 10.1371/journal.pone.0071393

60 Altgilbers, S., Dierks, C., Klein, S., Weigend, S. & Kues, W. A. Quantitative analysis of CRISPR/Cas9-mediated provirus deletion in blue egg layer chicken PGCs by digital PCR. Sci Rep 12, 15587 (2022). 10.1038/s41598-022-19861-7

61 Chen, L. et al. Dynamic 3D morphology of chick embryos and allantois depicted nondestructively by 3.0T clinical magnetic resonance imaging. Poult Sci 102, 102902 (2023). 10.1016/j.psj.2023.102902

62 Baggott, G. K. Development of Extra-embryonic Membranes and Fluid Compartments. Avian Biology Research 2, 21–26 (2009).

63 Simkiss, K. Water and Ionic Fluxes inside the Egg. American Zoologist 20, 385–393 (1980).

64 Everaert, N., Willemsen, H., Willems, E., Franssens, L. & Decuypere, E. Acid-base regulation during embryonic development in amniotes, with particular reference to birds. Respir Physiol Neurobiol 178, 118–128 (2011). 10.1016/j.resp.2011.04.023

65 Rideout, B. A. Investigating embryo deaths and hatching failure. Vet Clin North Am Exot Anim Pract 15, 155–162 (2012). 10.1016/j.cvex.2012.02.005

66 Romanoff, A. L. Critical Periods and Causes of Death in Avian Embryonic Development. The Auk 66, 264–270 (1949). 10.2307/4080357

67 Martinez-Rodero, I. et al. Blastocoel fluid aspiration improves vitrification outcomes and produces similar sexing results of in vitro-produced cattle embryos compared to microblade biopsy. Theriogenology 218, 142–152 (2024). 10.1016/j.theriogenology.2024.01.042

68 Powers, L. & Huntersville, N. (2021).

69 Zendehboudi, M. & Vesal, N. Comparison of cardiopulmonary effects of propofol, ketamine-propofol and isoflurane anesthesia in the domestic chicken (Gallus gallus domesticus). Vet Anaesth Analg 51, 449–457 (2024). 10.1016/j.vaa.2024.06.005

70 Horr, M., Sommerfeld, S., Silva, M. V. & Fonseca, B. B. A fast and simple protocol to anaesthesia in chicken embryos. Exp Anim 72, 294–301 (2023). 10.1538/expanim.22-0133

71 Zumbrink, L., Brenig, B., Foerster, A., Hurlin, J. & Wenzlawowicz, M. v. Electrical anaesthesia of male chicken embryos in the second third of the incubation period in compliance with animal welfare. European Poultry Science 84, 1–11 (2020). 10.1399/eps.2020.315

72 Aleksandrowicz, E. & Herr, I. Ethical euthanasia and short-term anesthesia of the chick embryo. ALTEX 32, 143–147 (2015). 10.14573/altex.1410031

73 Hatt, J. M., Kreyenbuhl, K. & Kummrow, M. [Methods of analgesia and euthanasia in backyard poultry]. Schweiz Arch Tierheilkd 165, 503–511 (2023). 10.17236/sat00398

74 Schlegel, L., Kleine, A. S., Doherr, M. G. & Fischer-Tenhagen, C. How to see stress in chickens: On the way to a Stressed Chicken Scale. Poult Sci 103, 103875 (2024). 10.1016/j.psj.2024.103875

