## Supplementary material for "*In Ovo* Sexing and Genotyping using PCR techniques: A Contribution to the 3R Principles in Chicken Breeding"

**Table S1** DNA Sexing and Genotyping Methods (Pilot Study, Phase I and II Studies)

| Genotyping Method | Sex-Specific Genomic Sequence | Variant | Primer Sequence<br>(5' - 3') | Amplicon Size<br>(bp) |
|---|---|---|---|---|
| Kompetitive allele-specific PCR (KASP) | Z: NC_052572.1:g.51980340-51980385<br>W: NC_052571.1:g.5554074-5554119 (reverse) | NC_052572.1:g. 51980363G > NC_052571.1:g. 5554096A (reverse) | FZ: CTTGACAAGCTACTGATTCGTCTG*<br>FW: CTTGACAAGTTACTGATTCGTCTA*<br>R: GWACTCTGTTGCCACGTTC[1] | 46 |
| multiplex PCR,<br>gel electrophoresis | Z: NC_ 052572.1:g.51980348-51980940<br>W: NC_052571.1:g.5554111-5553685 (reverse) | sex-specific amplicon size: CHD1-Z and CHD1-W gene | F1: GCTACTGATTCGTCTGCGAGA*<br>F2: GTTACTGATTCGTCTACGAGA**<br>R: ATTGAAATGATCCAGTGCTTG** | 593 (Z)<br>447 (W) |
| Kompetitive allele-specific PCR (KASP) | NC_052532.1:g.65226429-65226505 (wild-type)<br>KC632578.1:g.78-157 (reverse) (Araucana-specific) | blue egg shell :<br>EAV-HP insertion upstream of SLCO1B3 | F1: TGACCAGCGTAGATAAACATG<br>F2: AACACACCTCGTTTCCCTC<br>R: AGCAGTTTTAATCTTGTCTCCTC | 77/80 |
| multiplex PCR, gel electrophoresis | NC_052532.1:g. 65226368-65226730 (wild-type)<br>NC_052532.1:g.65226368-65226369 + KC632578.1:g.4293-4457 (reverse) (Araucana-specific) | blue egg shell :<br>EAV-HP insertion upstream of SLCO1B3 | F: GCATTTCACAAACGGGTGTA***<br>R1: AAAACCACAAAGGTAATGTTCA***<br>R2: CCCAGCAGTAAGCCCTACAT*** | 363/167 |

*Reference: Dierks *et al.*, 2022[1], **Fridolfsson *et al.*, 1999[2]; ***Reference: Wragg *et al.*, 2013[3]

**Table S2** Sexing and EAV-HP-Insertion Genotyping Success Rates (Pilot study)

| Incubation length (h) | Number of Eggs | DNA Detectable on Agarose Gel** | Mean DNA Concentration (ng/µl) | Successful Sexing (PCR) | Successful Sexing (KASP) | Discrepancies Tissue vs. Allantois (Sexing) | Successful Genotyping (PCR) | Successful Genotyping (KASP) | Discrepancies Tissue vs. Allantois (Genotyping) | Macroscopic Sex Verification (day 10) |
|---|---|---|---|---|---|---|---|---|---|---|
| 96 | 30 | 90% | 65.9 | 80% | 70% | 0 | 67% | 73% | 0 | N/A[#] |
| 144 | 30 | 100% | 75.9 | 100% | 97% | 2 | 93% | 93% | 2 | N/A[#] |
| 192 | 30 | 100% | 72.1 | 100% | 100% | 0 | 100% | 100% | 0 | N/A[#] |
| 240 | 30 | 100% | 75.8 | 100% | 100% | 0 | 100% | 97% | 0 | 30 |

**See related Fig. S6; [#]Inspection of the gonads after opening of the abdominal cavity. Sexual gonadal dimorphism is difficult to assess visually at these early embryonic stages. The gonads remain bipotential until HH 29-31, exhibiting no visible left-right asymmetry[4] (see related Fig. S2); KASP: Kompetitive Allele Specific PCR

**Table S3** Hatching Results: Comparison of Hatch Rates between *In Ovo* Sexed Chicks (sum of ED4-ED7) and Untreated Eggs in Phase II Study

| Chicken Line | | Age of flock (weeks) | Kind of hatch | Eggs total | Eggs fertilized | Fertilized Eggs Punctured *in ovo* | Unfertilized[a] | Vent Sexing | Hatched N | Hatchability[c] |
|---|---|---|---|---|---|---|---|---|---|---|
| Brown layer N = 447 | | 63 | single | 240 | **240** | 240 | 0% | female | 81 | 47% |
| | | | | | | | | male | 93 | 53% |
| | | | | | | | | total | **174** | **73%** |
| | | | flock control | 207 | **153** | 26[b] | 26% | female | 46 | 48% |
| | | | | | | | | male | 50 | 52% |
| | | | | | | | | total | **96** | **63%** |
| Control groups | N = Sum of 3 hatches | 63 | flock | N/A | N/A | N/A | 13-16% | female | N/A | 37-41% |
| White layer N = 449 | | 42 | single | 239 | **239** | 239 | 0% | female | 111 | 50% |
| | | | | | | | | male | 109 | 50% |
| | | | | | | | | total | **220** | **92%** |
| | | | flock control | 210 | **197** | 22[b] | 6% | female | 84 | 46% |
| | | | | | | | | male | 98 | 54% |
| | | | | | | | | total | **182** | **92%** |
| Control groups | N = Sum of 5 hatches | 42 | flock | N/A | N/A | N/A | 2-7% | female | N/A | 44-47% |

[a]In the single-hatched and *in ovo* sexed groups unfertilized eggs were excluded prior to hatching. In the flock control, unfertilized eggs were returned into the incubator. [b]A subset of flock control eggs (fertilized) were punctured for testing *in ovo* sampling procedure and were not excluded. [c]Total hatchability was calculated based on the total number of fertilized eggs. Male and female hatch rates were calculated based on the total number of hatched chicks.

**Table S4** Sexing and iCaspase9-GFP Genotyping (Phase III Study)

| Trial* | Incubation length (h) | Number of Eggs | Successful Sexing (PCR) | Discrepancies Tissue vs. Allantois (Sexing) | Successful Genotyping (PCR) | Discrepancies Tissue vs. Allantois (Genotyping) | Survival Rate Post-Puncture (Day 11) | Male vs. Female Embryos (n) | Genotype of Embryos (n) (hom:het:wt)** |
|---|---|---|---|---|---|---|---|---|---|
| 1 | 190 | 34 | 100% | 0 | 100% | 0 | 94% | 17:17 | 11:18:5 |
| 2 | 190 | 33 | 100% | 0 | 100% | 1 | 88% | 15:18 | 5:21:6 |
| 3 | 190 | 33 | 100% | 1 | 100% | 0 | 94% | 19:13 | 7:21:5 |

*The experiment was repeated weekly. **based on concordant sex and genotype identification between allantoic fluid and tissue samples; hom: homozygous, he: heterozygous iCaspase9, wt: wildtype (iCaspase9 negative)

**Table S5** Sexing and iCaspase9-GFP Genotyping (Phase III Study, 2$^{nd}$ experimental phase)

| Category | Number of eggs incubated | Unfertilized eggs | Early losses (<ED7) | Fertilized eggs (ED7) | Samples yielding a PCR result | Samples with no initial PCR result | Discrepancies tissue vs. AF (Sexing & Genotyping) |
|---|---|---|---|---|---|---|---|
| Number (n) | 177 | 33 | 3 | 141 | 138 | 19 | 3 |
| Percentage (%) | 100% | 18.6% | 1.7% | 79.7% | 97.9% (of 141) | 13.5% (of 141) | 0.7% (of 141) |

AF=Allantoic fluid, control=embryonic tissue

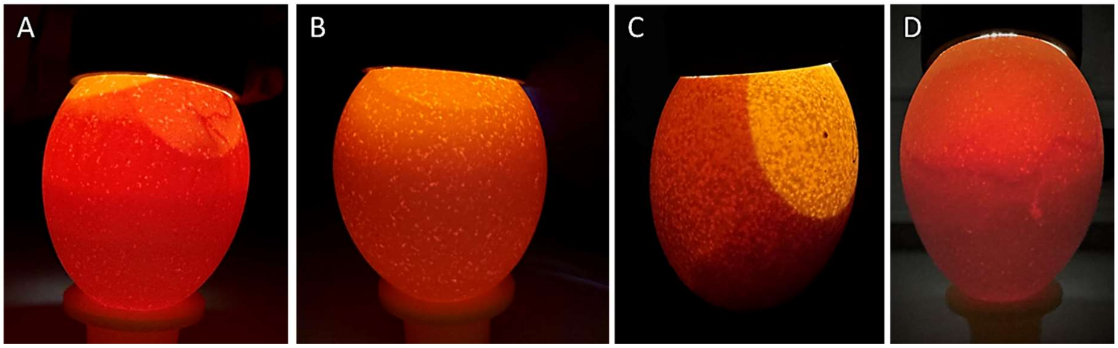

**Fig. S1 Egg Candling Inspection**

Brown eggs were candled at ED7 using an LED light source directed from the blunt end through the air chamber. **(A)** Fertilized egg with a visible air chamber at the blunt end and a clearly defined liquid-filled area (amnion) where blood vessels of the chorioallantoic membrane (CAM) are evident and embryonic movement can be observed. **(B)** Unfertilized egg with uniform, bright translucency throughout. **(C)** Displaced air chamber away from the blunt end can be caused by factors such as rough handling, aging of the egg, or improper storage conditions (incorrect temperature and/or humidity). **(D)** A blood ring, indicating embryonic death or failed development.

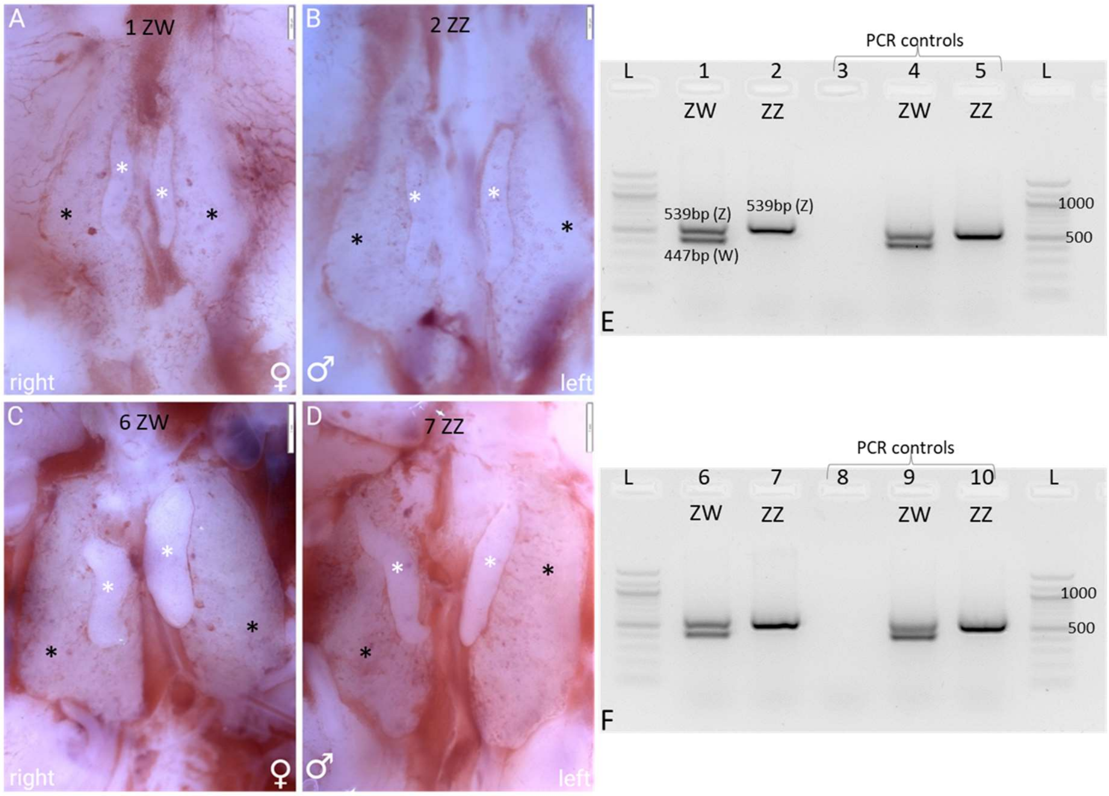

**Fig. S2 Left Right Asymmetry in Chicken Gonads at Embryonic Day 6.5 (ED6.5) versus ED10**

**A-D** Comparison of chicken embryonic gonads at ED6.5 and ED10 of embryonic development, visualized by light microscopy (Olympus Stereomicroscope SZ61 with Olympus EP50 camera, EVIDENT Europe GmbH, Hamburg, Germany). The gonads are indicated by white asterisks. The gonads are located on the medioventral surface of the mesonephric kidney (black asterisks). **(A-B)** Chick embryo gonads at ED6.5 display no sexual dimorphism. Scale bar = 500 µm. **(C-D)** Chick embryo gonads at ED10 display sexual dimorphism. Scale bar = 1 mm. **(D)** Males possess two similarly sized, elongated testes, while **(C)** females exhibit a prominent, bean-shaped left ovary that is wider in its transverse dimension than the testes, and a regressing right ovary. **(E-F)** Sexing PCR was performed on the gonadal tissue corresponding to the photos shown in (A-D). **(E)** L=100 bp ladder, 1=A (female), 2=B (male), 3=$H_20$, negative control, 4=female control, 5=male control. **(F)** L=100 bp ladder, 6=C (female), 7=D (male), 8=$H_20$, negative control, 9=female control, 10=male control.

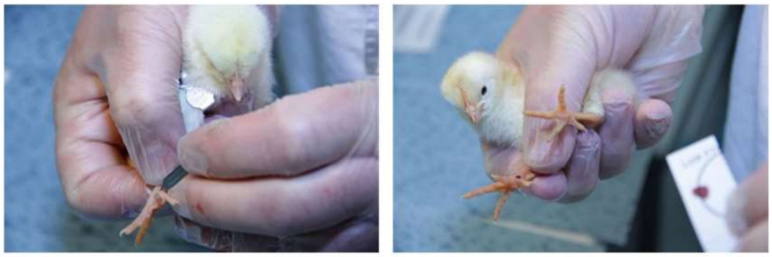

**Fig. S3 Capillary Blood Sampling from the Foot of a Chick**[5]

**(A)** Blood sampling was performed by a short puncture of the Vena metatarsalis plantaris superficialis medialis with a lancet. **(B)** The small drop of blood was collected on medium filtration capacity filter paper cards[6] (FTA® card, Schleicher & Schuell GmbH, Whatman, Inc., Maidstone, U.K.). This blood volume was sufficient for subsequent DNA extraction, as chicken erythrocytes are nucleated. After puncture, gentle pressure was applied to the wound with a soft pad until bleeding stopped; hemostasis occurred rapidly.

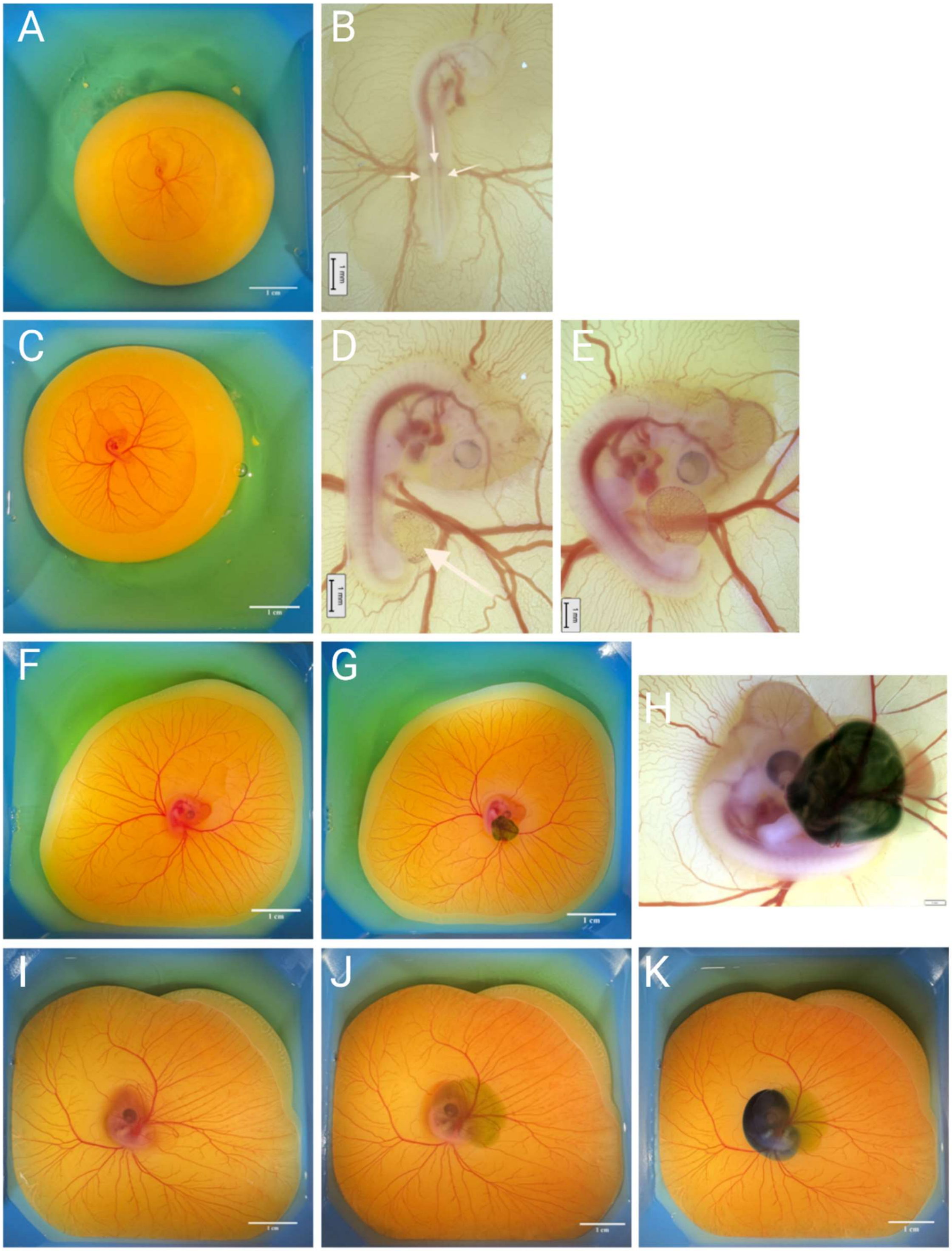
A
B
1 cm
1 mm
C
D
E
1 cm
1 mm
1 mm
F
G
H
1 cm
1 cm
I
J
K
1 cm
1 cm
1 cm

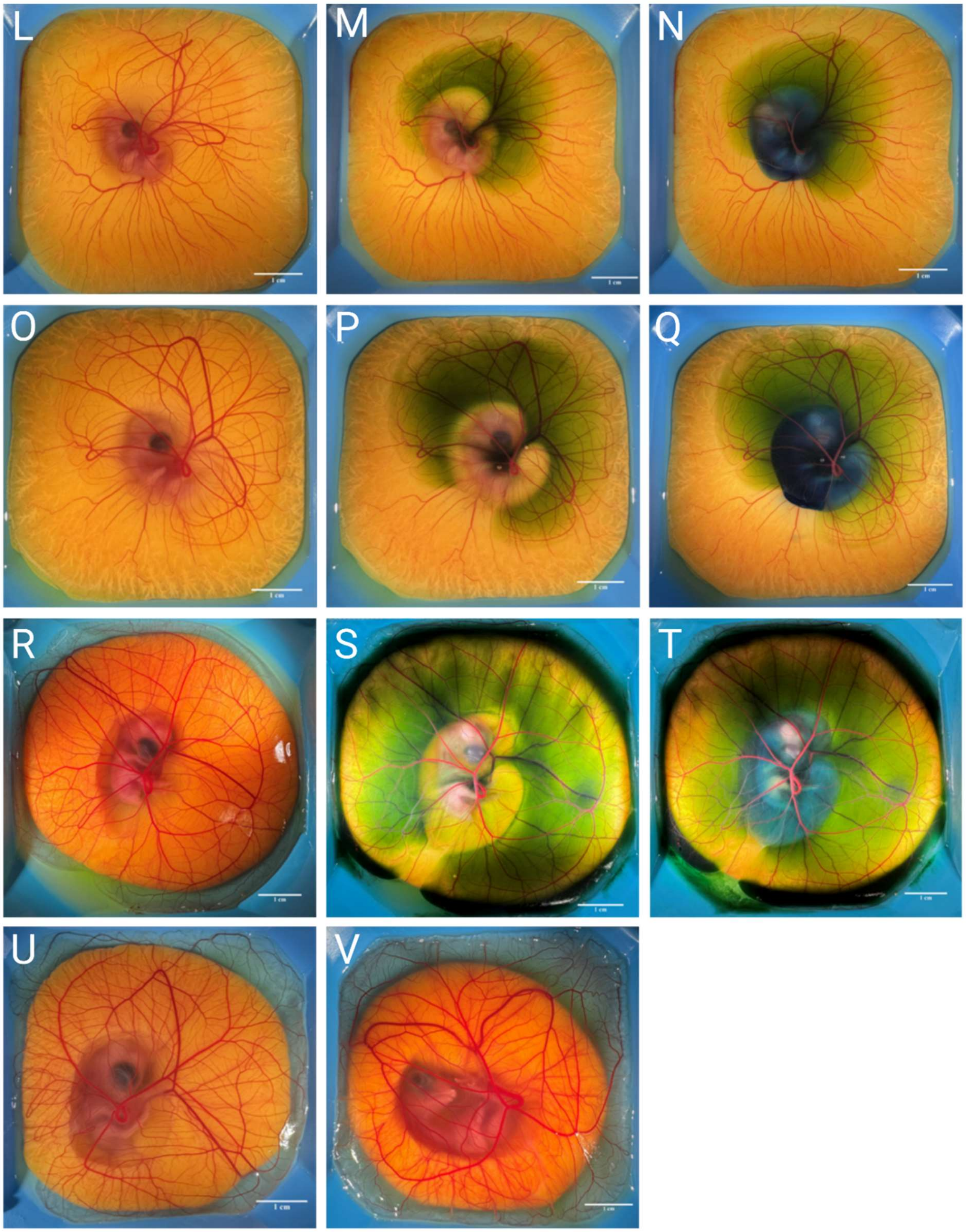

**Fig. S4** ***Ex ovo*** **Culture of Chicken Embryos (Embryonic day 2 to 11 (ED2-ED11))**

Illustration of the spatiotemporal development of the chorioallantoic membrane (CAM) and embryonic development within the amnion cavity in a shell-less culture system. Dyes (green and blue) were used to visualize the CAM and amnion, respectively. **(A)** Photograph of a 2.5-day incubated chicken embryo. **(B)** A microscopic view of (A) depicting the embryo at Hamburger and Hamilton[7] stage 16 (HH16): The amnion remains open (white arrows), the allantoic vesicle has not yet formed; scale bar: 1 mm. **(C-D)** Early embryo after 75 hours of incubation. **(D)** The amnion is closed and the allantois is visible as a small vesicle, representing an evagination from the endodermal hindgut (white arrow); scale bar: 1 mm. **(E)** Embryo at 96 hours of incubation. The dorsal contour, from hindbrain to tail, exhibits a curved line. The allantois is comparable in size to the midbrain; scale bar: 1 mm. The figure further details the developmental progression with representative photographs at the following incubation times: **(F-H)** 116 hours **(I-K)** 144 hours **(L-N)** 168 hours **(O-Q)** 192 hours **(R-T)** 240 hours, **(U)** 264 hours and **(V)** 280 hours. For each stage, the corresponding panels illustrate: **(F, I, L, O, R, U, V)** the overall morphology, **(G, H, J, M, P, S)** the green-stained allantoic cavity, and **(K, N, Q, T)** the blue-stained amniotic cavity. **(H)** Microscopic view, magnified from **(G);** The amnion, a transparent sac surrounding the embryo, contains limited fluid at this stage of development, making staining difficult without membrane disruption. Scale bar: 1 cm unless otherwise stated.

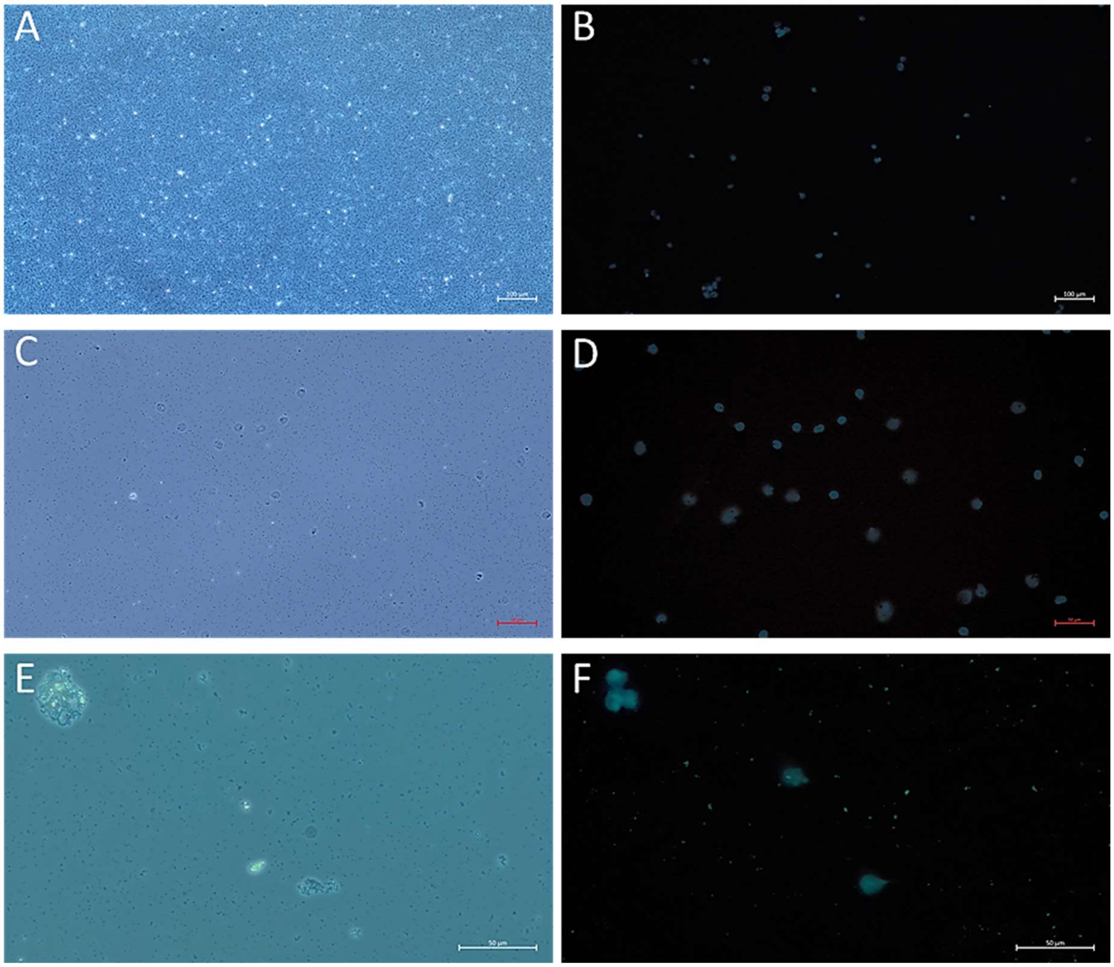

**Fig. S5 Cell Detection in Fluid Samples (Hoechst Staining)**

**(A)** 10x magnification: Microscopic view of *in ovo* fluid collected at ED4. The background exhibits abundant unstained sediment. The pellet formed after centrifugation of this fluid appears white, dense, and readily visible. **(B)** 10x magnification: Hoechst staining of nuclei in the field of view shown in (A). **(C)** 20x magnification: Allantoic fluid collected at ED7. The background is clearer, with less visible sediment after centrifugation. **(D)** 20x magnification: Hoechst staining of nuclei in the field of view shown in (C). **(E)** 40x magnification: Allantoic fluid collected at ED7, showing cell clusters surrounded by undefined debris or sediment. **(F)** 40x magnification: Hoechst staining of the field of view shown in (E), revealing smaller and larger cells with blue nuclear staining. Scale bars: 50 µm.

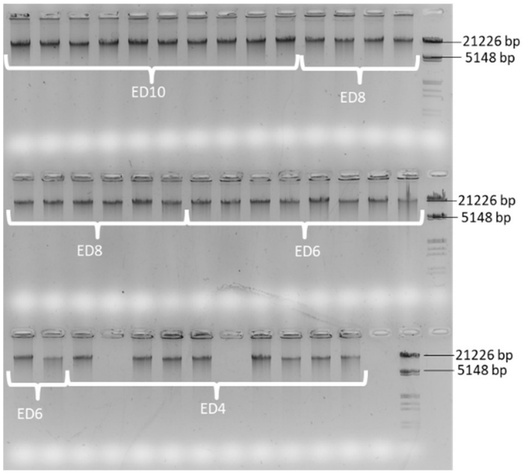

**Fig. S6 Evaluation of DNA Integrity from Allantoic Fluid Samples at different Developmental Stages**
This figure displays one representative gel from the Pilot study evaluating DNA quality control on a 0.8 % agarose. Two microliters of each of 10 samples per isolation day (ED4, ED6, ED8, ED10) were loaded onto the gel. The Roche DNA Molecular Weight Marker III was used as a size standard. The highest amounts of DNA were observed in samples isolated at ED10. Two samples from ED4 isolation failed to show detectable DNA. The Repli-g-Minikit from Qiagen (Whole Genome Amplification Kit) yielded high-quality, high-molecular-weight DNA.

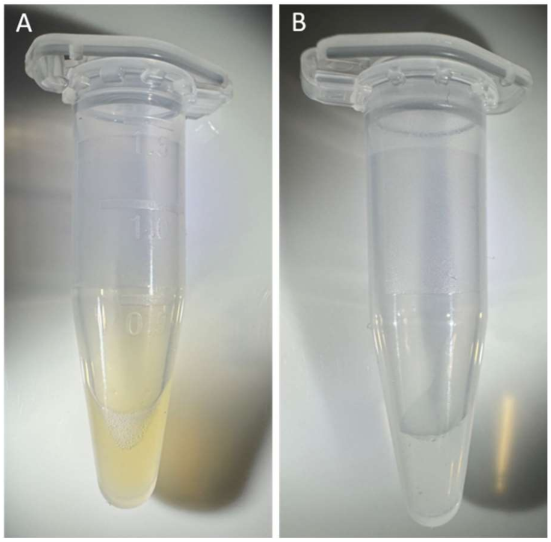

**Fig. S7 Visual Comparison of *in ovo* and Allantoic Fluid Coloration**
**(A)** Yellowish fluid extracted from embryos at ED4 to ED6 (*in ovo* fluid). **(B)** Clear fluid extracted from embryos at ED7 and beyond (most likely allantoic fluid).

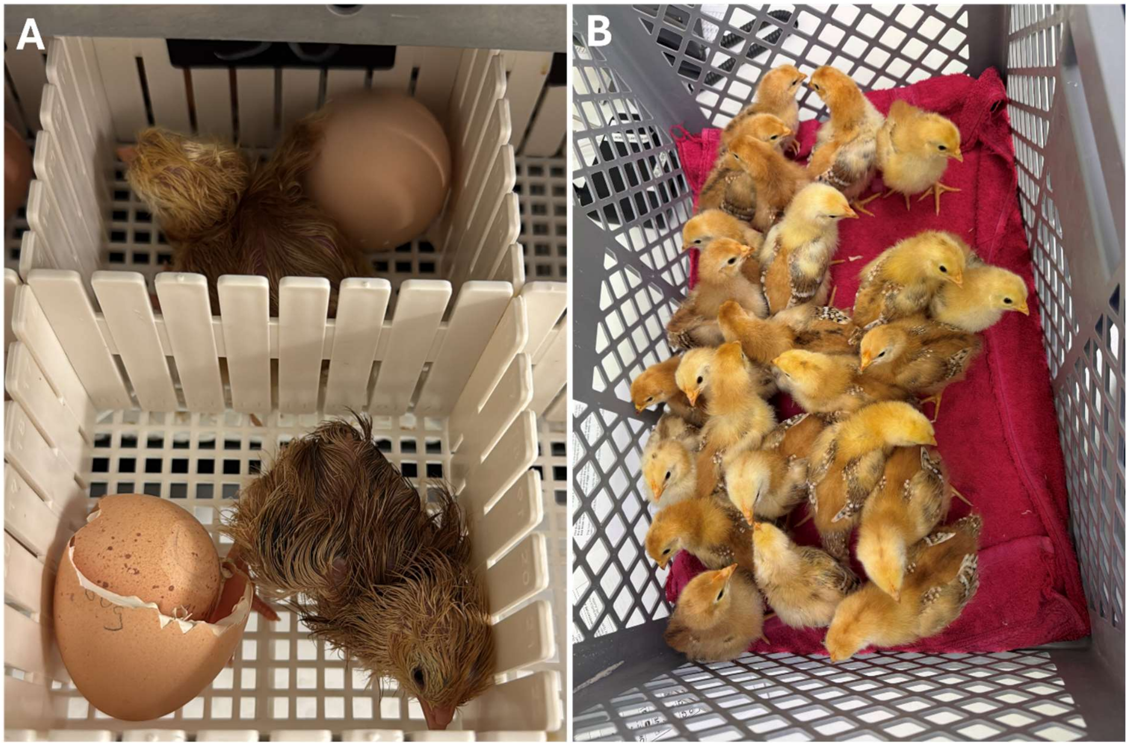

**Fig.S8 iCaspase9 Chicks (*in ovo* sexed and genotyped)**

**A)** Newly hatched chicks in individual compartments within the hatcher. The chorioallantois membrane (CAM) from their respective eggs serves as a reliable source for DNA extraction, enabling accurate sexing and genotyping without invasive procedures. **B)** 24 healthy, 10-day-old chicks (iCaspase9/L68).

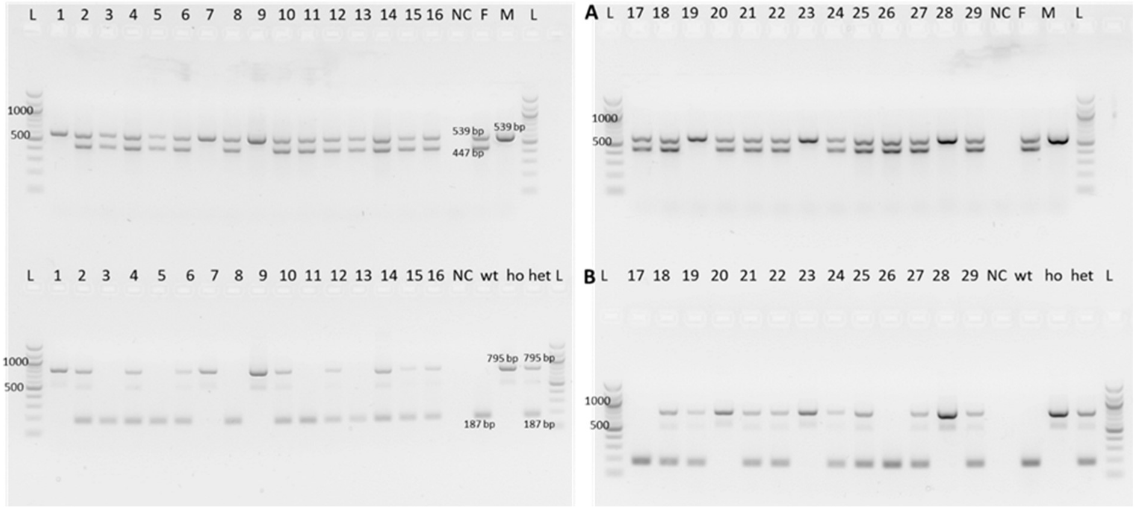

**Fig. S9 PCR-Based Sexing and Genotyping of Chorioallantoic Membrane (CAM) Samples (Phase Study III)**
Molecular analysis of 29 CAM samples post hatch, confirming the (**A**) *in ovo* sexing and corresponding (**B**) genotyping (iCaspase9)[8] results previously obtained at ED8. The samples correspond to the 29 eggs chosen for hatching. N: $H_20$, F: female control, M: male control, wt: wildtype (iCaspase9 negative; 187 bp), hom: homozygous iCaspase9 (795bp), het: heterozygous iCaspase9 (795bp+187 bp). L: 100bp ladder.

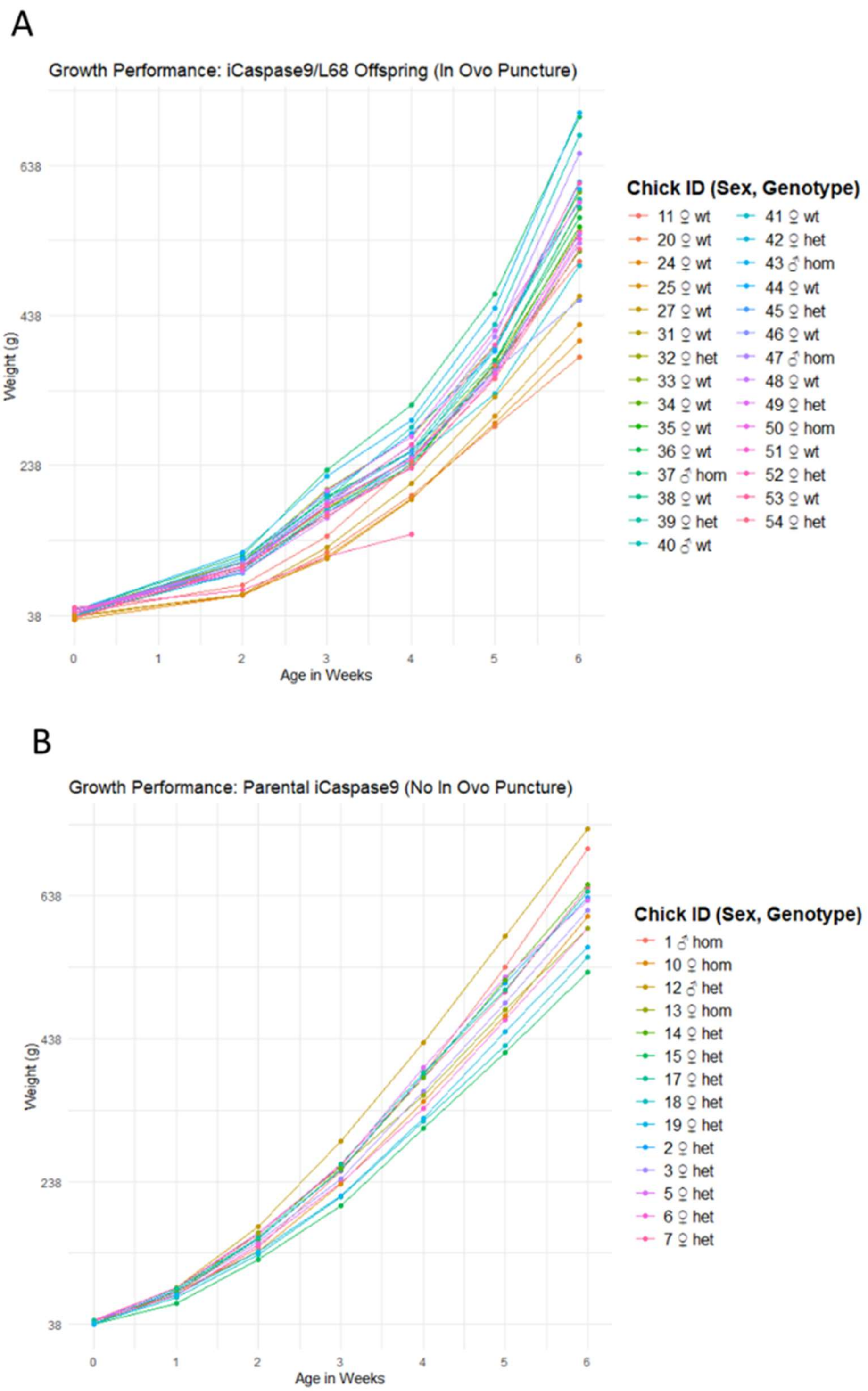

**Fig. S10 Growth Performance iCaspase9 Offspring (*In Ovo* Puncture) vs. Parental (No *In Ovo* Puncture)**
Comparison of weekly body weight (g) over six weeks between (A) iCaspase9/L68 offspring (subjected to *in ovo* puncture, 24 chicks), and (B) parental iCaspase9 controls (no *in ovo* Puncture, 14 chicks). Each colored line represents the growth curve of an individual chick. Sex: (♂=male, ♀=female; genotype: wt=wild-type (iCaspase9 negative), het=heterozygous iCaspase9, hom=homozygous iCaspase9.

1 Dierks, C., Altgilbers, S., Weigend, A., Preisinger, R. & Weigend, S. Sexing assay for chickens and other birds for large-scale application based on a conserved sequence variant in CHD1 genes on W and Z chromosomes. *Anim Genet* **53**, 235-237 (2022). https://doi.org:10.1111/age.13176

2 Fridolfsson, A. K. & Ellegren, H. A simple and universal method for molecular sexing of non-ratite birds. *J. Avian Biol.* **30**, 116-121 (1999).

3 Wragg, D. *et al.* Endogenous retrovirus EAV-HP linked to blue egg phenotype in Mapuche fowl. *PLoS One* **8**, e71393 (2013). https://doi.org:10.1371/journal.pone.0071393

4 Cutting, A. D., Bannister, S. C., Doran, T. J., Sinclair, A. H., Tizard, M. V. & Smith, C. A. The potential role of microRNAs in regulating gonadal sex differentiation in the chicken embryo. *Chromosome Res* **20**, 201-213 (2012). https://doi.org:10.1007/s10577-011-9263-y

5 Altgilbers, S. *Etablierung des Gene Editing beim Huhn anhand der Modifikation des SLCO1B3-Gens*, Tierärztliche Hochschule Hannover, (2020).

6 Suriyaphol, G., Kunnasut, N., Sirisawadi, S., Wanasawaeng, W. & Dhitavat, S. Evaluation of dried blood spot collection paper blotters for avian sexing by direct PCR. *Br. Poult. Sci.* **55**, 321-328 (2014). https://doi.org:10.1080/00071668.2014.925087

7 Hamburger & Hamilton. A series of normal stages in the development of the chick embryo. *J Morphol 88(1): 49-92.* (1951).

8 Ballantyne, M. *et al.* Direct allele introgression into pure chicken breeds using Sire Dam Surrogate (SDS) mating. *Nat Commun* **12**, 659 (2021). https://doi.org:10.1038/s41467-020-20812-x
